# Emergent rules for codon choice elucidated by editing rare arginine codons in *Escherichia coli*

**DOI:** 10.1101/069385

**Authors:** Michael G. Napolitano, Matthieu Landon, Christopher J. Gregg, Marc J. Lajoie, Lakshmi N. Govindarajan, Joshua A. Mosberg, Gleb Kuznetsov, Daniel B. Goodman, Oscar Vargas-Rodriguez, Farren J. Isaacs, Dieter Söll, George M. Church

## Abstract

The degeneracy of the genetic code allows nucleic acids to encode amino acid identity as well as non-coding information for gene regulation and genome maintenance. The rare arginine codons AGA and AGG (AGR) present a case study in codon choice, with AGRs encoding important transcriptional and translational properties distinct from the other synonymous alternatives (CGN). We created a strain of *Escherichia coli* with all 123 instances of AGR codons removed from all essential genes. We readily replaced 110 AGR codons with the synonymous CGU, but the remaining thirteen “recalcitrant” AGRs required diversification to identify viable alternatives. Successful replacement codons tended to conserve local ribosomal binding site-like motifs and local mRNA secondary structure, sometimes at the expense of amino acid identity. Based on these observations, we empirically defined metrics for a multi-dimensional “safe replacement zone” (SRZ) within which alternative codons are more likely to be viable. To further evaluate synonymous and non-synonymous alternatives to essential AGRs, we implemented a CRISPR/Cas9-based method to deplete a diversified population of a wild type allele, allowing us to exhaustively evaluate the fitness impact of all 64 codon alternatives. Using this method, we confirmed relevance of the SRZ by tracking codon fitness over time in 14 different genes, finding that codons that fall outside the SRZ are rapidly depleted from a growing population. Our unbiased and systematic strategy for identifying unpredicted design flaws in synthetic genomes and for elucidating rules governing codon choice will be crucial for designing genomes exhibiting radically altered genetic codes.

**Significance Statement:** This work presents the genome-wide replacement of all rare AGR arginine codons in the essential genes of *Escherichia coli* with synonymous CGN alternatives. Synonymous codon substitutions can lethally impact non-coding function by disrupting mRNA secondary structure and ribosomal binding site-like motifs. Here we quantitatively define the range of tolerable deviation in these metrics and use this relationship to provide critical insight into codon choice in recoded genomes. This work demonstrates that genome-wide removal of AGR is likely to be possible, and provides a framework for designing genomes with radically altered genetic codes.

## Main Text

The genetic code possesses inherent redundancy (1), with up to six different codons specifying a single amino acid. While it is tempting to approximate synonymous codons as equivalent (2), most prokaryotes and many eukaryotes (3, 4) display a strong preference for certain codons over synonymous alternatives (5, 6). While different species have evolved to prefer different codons, codon bias is largely consistent within each species (5). However, within a given genome, codon bias differs among individual genes according to codon position, suggesting that codon choice has functional consequences. For example, rare codons are enriched at the beginning of essential genes (7, 8), and codon usage strongly affects protein levels (9-11), especially at the N-terminus (12). This suggests that codon usage plays a poorly understood role in regulating protein expression. Several hypotheses attempt to explain how codon usage mediates this effect, including but not limited to: facilitating ribosomal pausing early in translation to optimize protein folding (13); adjusting mRNA secondary structure to optimize translation initiation or to modulate mRNA degradation; preventing ribosome stalling by co-evolving with tRNA levels (6); providing a “translational ramp” for proper ribosome spacing and effective translation (14); and providing a layer of translational regulation for independent control of each gene in an operon (15). Additionally, codon usage may impact translational fidelity (16), and the proteome may be tuned by fine control of the decoding tRNA pools (17). Although Quax et al. provide an excellent review of how biology chooses codons, systematic and exhaustive studies of codon choice in whole genomes are lacking (18). Studies have only begun to empirically probe the effects of codon choice in a relatively small number of reporter genes (12, 19-22). Several important questions must be answered as a first step towards designing custom genomes exhibiting new functions—How flexible is genome-wide codon choice? How does codon choice interact with the maintenance of cellular homeostasis? What heuristics can be used to predict which codons will conserve genome function?

Replacing all essential instances of a codon in a single strain would provide valuable insight into the constraints that determine codon choice and aid in the design of recoded genomes. Although the UAG stop codon has been completely removed from *Escherichia coli* (23), no genome-wide replacement of a sense codon has been reported. While the translation function of the AGG codon has been shown to permit efficient suppression with nonstandard amino acids (24-26), AGG necessarily remains translated as Arg in each of these studies. No study has yet demonstrated that all AGR codons (or all instances of any sense codon) can be removed from essential genes, nor explained why certain AGR codons could not be changed successfully. These insights are crucial for unambiguously reassigning AGR translation function.

We chose to study the rare arginine codons AGA and AGG (termed AGR according to IUPAC conventions) because the literature suggests that they are among the most difficult codons to replace and that their similarity to ribosome binding sequences underlies important non-coding functions (8, 27-30). Furthermore, their sparse usage (123 instances in the essential genes of *E. coli* MG1655 and 4228 instances in the entire genome (Table 1, S1)) made replacing all AGR instances in essential genes a tractable goal, with essential genes serving as a stringent test set for identifying any fitness impact from codon replacement (31). Additionally, recent work has shown the difficulty of directly mutating some AGR codons to other synonymous codons (25), although the authors do not explain the mechanism of failure or report successful implementation of alternative designs. We attempted to remove all 123 instances of AGR codons from essential genes by replacing them with the synonymous CGU codon. CGU was chosen to maximally disrupt the primary nucleic acid sequence (<u>A</u>G<u>R</u>→<u>C</u>G<u>U</u>). We hypothesized that this strategy would maximize design flaws, thereby revealing rules for designing genomes with reassigned genetic codes. Importantly, individual codon targets were not inspected *a priori* in order to ensure an unbiased empirical search for design flaws.

**Table 1.**
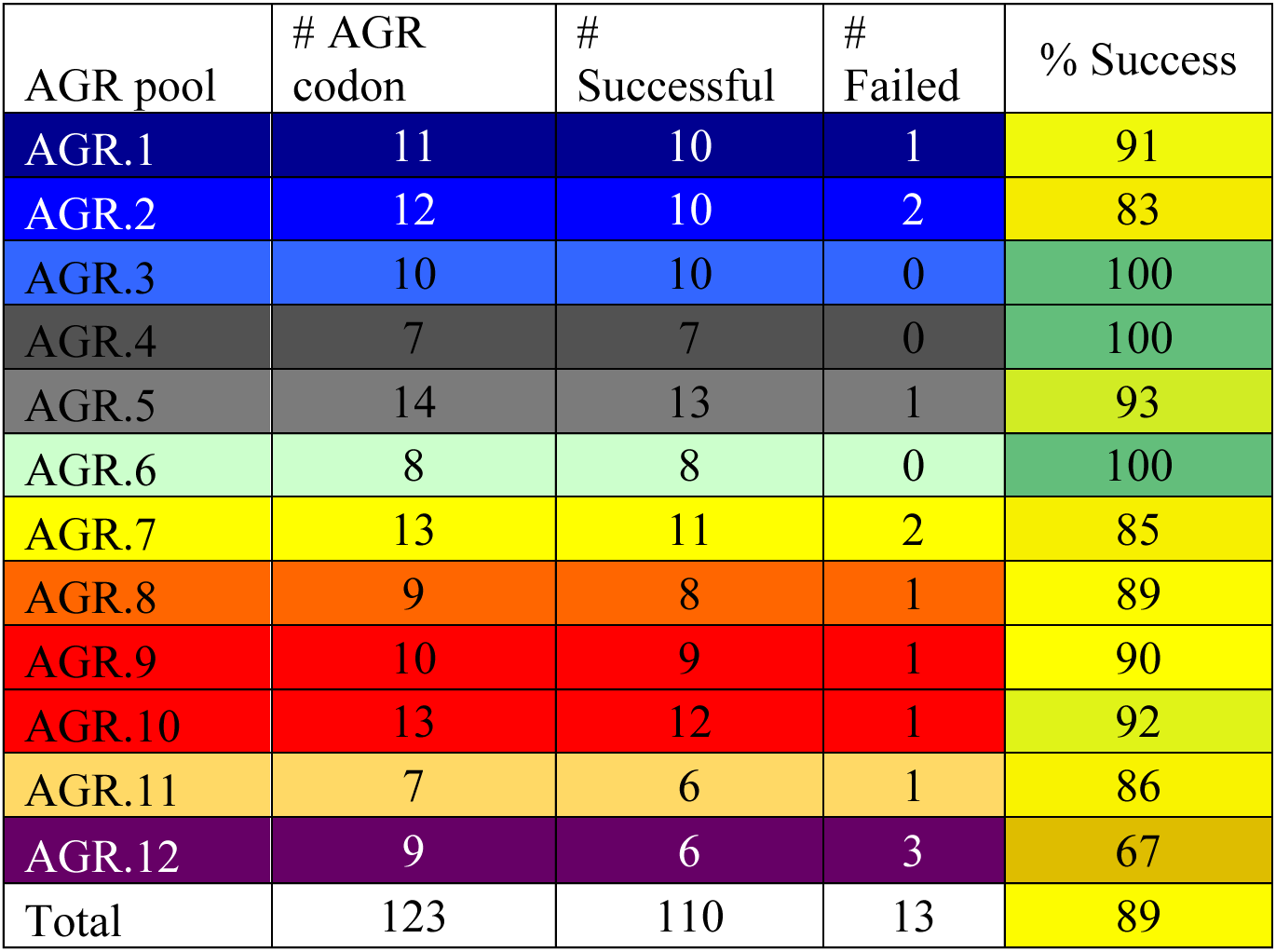
Summary of AGR codons changed by location in the genome, and failure rates by pool.

To construct this modified genome, we used co-selection multiplex automatable genome engineering (CoS-MAGE) (32, 33) to create an *E. coli* strain (C123) with all 123 AGR codons removed from its essential genes (Figure 1A and see Table S1 for a complete list of AGR codons in essential genes). CoS-MAGE leverages lambda red-mediated recombination (34, 35) and exploits the linkage between a mutation in a selectable allele (e.g. *tolC*) to nearby edits of interest (e.g., AGR conversions), thereby enriching for cells with those edits (Figure S1). To streamline C123 construction, we chose to start with *E. coli* strain EcM2.1, which was previously optimized for efficient lambda red-mediated genome engineering (33, 36). Using CoS-MAGE on EcM2.1 improves allele replacement frequency by 10-fold over MAGE in non-optimized strains but performs optimally when all edits are on the same replichore and within 500 kilobases of the selectable allele (33). To accommodate this requirement, we divided the genome into 12 segments containing all 123 AGR codons in essential genes. A *tolC* cassette was moved around the genome to enable CoS-MAGE in each segment, allowing us to rapidly prototype each set of AGR→CGU mutations across large cell populations *in vivo*. (Please see the ‘General Replacement Strategy’ and ‘Troubleshooting Strategy’ sections of the Materials & Methods for a more detailed discussion). Of the 123 AGR codons in essential genes, 110 could be changed to CGU by this process (Figure 1), revealing considerable flexibility of codon usage for most essential genes. Allele replacement (in this case, AGR→CGU codon substitution) frequency varied widely across these 110 permissive codons, with no clear correlation between allele replacement frequency and normalized position of the AGR codon in a gene (Figure 2A).

**Figure 1.**
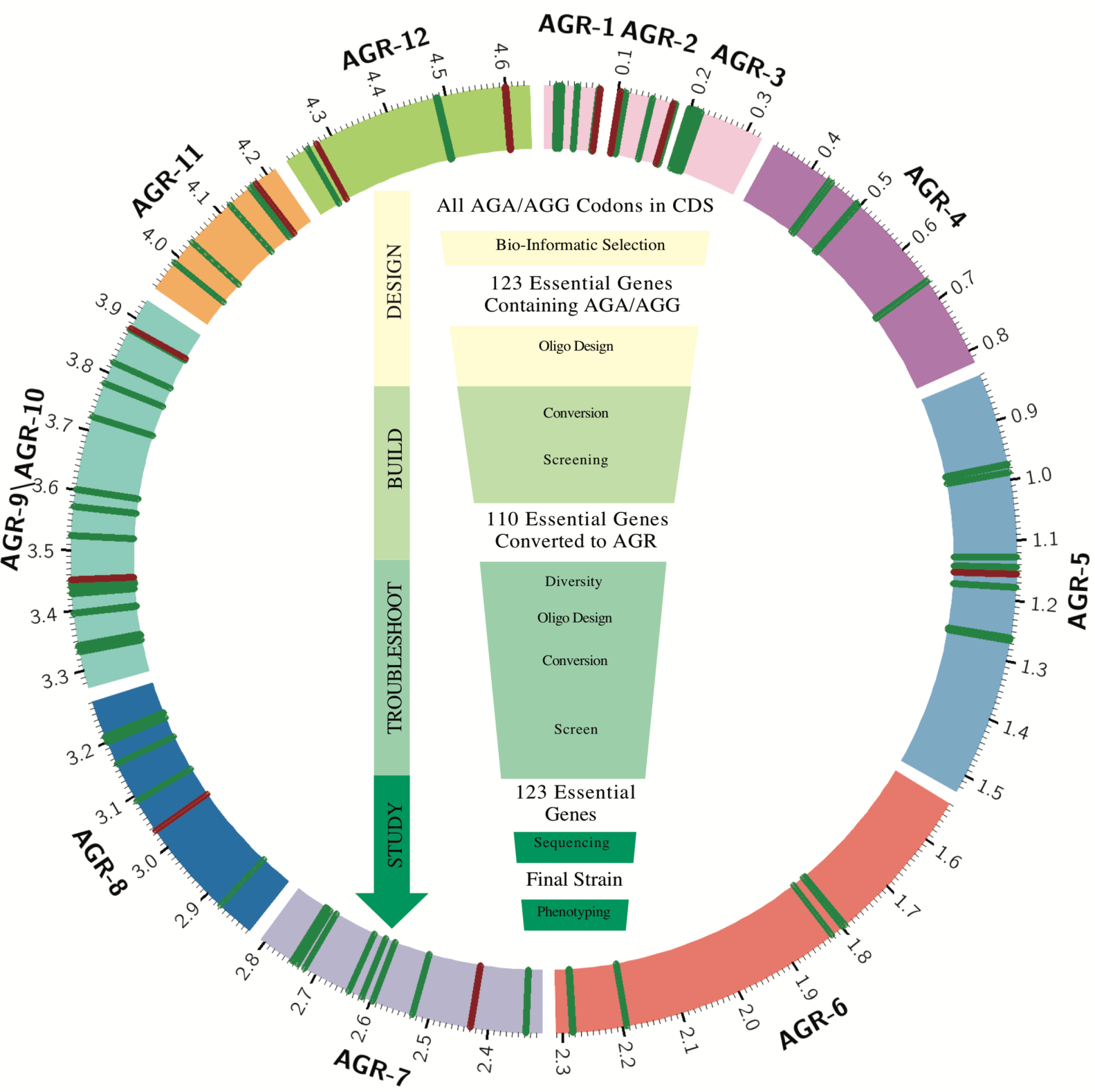
Construction of strain C123. (inner) Workflow used to create and analyze strain C123. The DESIGN phase involved identification of 123 AGR codons in the essential genes of *Escherichia coli*. MAGE oligos were designed to replace all instances of these AGR codons with the synonymous CGU codon. The BUILD phase used CoS-MAGE to convert 110 AGR codon to CGU and to identify 13 AGR codons that required additional troubleshooting. The *in vivo* TROUBLESHOOTING phase resolved the 13 codons that could not be readily converted to CGU and identified mechanisms potentially explaining why AGR→CGU was not successful. In the STUDY Phase, next-generation sequencing, evolution and phenotyping was performed on strain C123. **(outer)** Schematic of the C123 genome (Nucleotide 0 oriented up; numbering according to strain MG1655). Exterior labels indicate the set groupings of AGR codons. Successful AGR→CGU (110 instances) conversions are indicated by radial green lines, and recalcitrant AGR codons (13 instances) are indicated by radial red lines.

The remaining 13 AGR→CGU mutations were not observed, suggesting a codon substitution frequency of less than our detection limit of 1% of the bacterial population (Materials & Methods, Table S6). These “recalcitrant codons” were assumed to be deleterious or non-recombinogenic and were triaged into a troubleshooting pipeline for further analysis (Figure 1). Interestingly, all except for one of the thirteen recalcitrant codons were co-localized near the termini of their respective genes, suggesting the importance of codon choice at these positions — seven were at most 30 nt downstream of the start codon, while five were at most 30 nucleotides (nt) upstream of the stop codon (Figure 2A, lower panel, Table S8). Due to our unbiased design strategy, we anticipated that several AGR→CGU mutations would present obvious design flaws such as introducing non-synonymous mutations (two instances) or RBS disruptions (four instances) in overlapping genes. For example, *ftsI*_AGA1759 overlaps the second and third codons of *murE*, an essential gene, introducing a missense mutation (*murE* D3V) that may impair fitness. Replacing *ftsI*_AGA with <u>C</u>GA successfully replaced the forbidden AGA codon while conserving the primary amino acid sequence of MurE with a minimal impact on fitness (Figure 3A, Table S6). Similarly, *holB*_AGA4 overlaps the upstream essential gene *tmk*, and replacing AGA with CGU converts the *tmk* stop codon to Cys, adding 14 amino acids to the C-terminus of *tmk*. While some C-terminal extensions are well-tolerated in E. coli (37), extending *tmk* appears to be deleterious. We successfully replaced *holB*_AGA with <u>C</u>G<u>C</u> by inserting three nucleotides comprising a stop codon before the *holB* start codon. This reduced the *tmk/holB* overlap, and preserved the coding sequences of both genes (Figure S2A).

**Figure 2.**
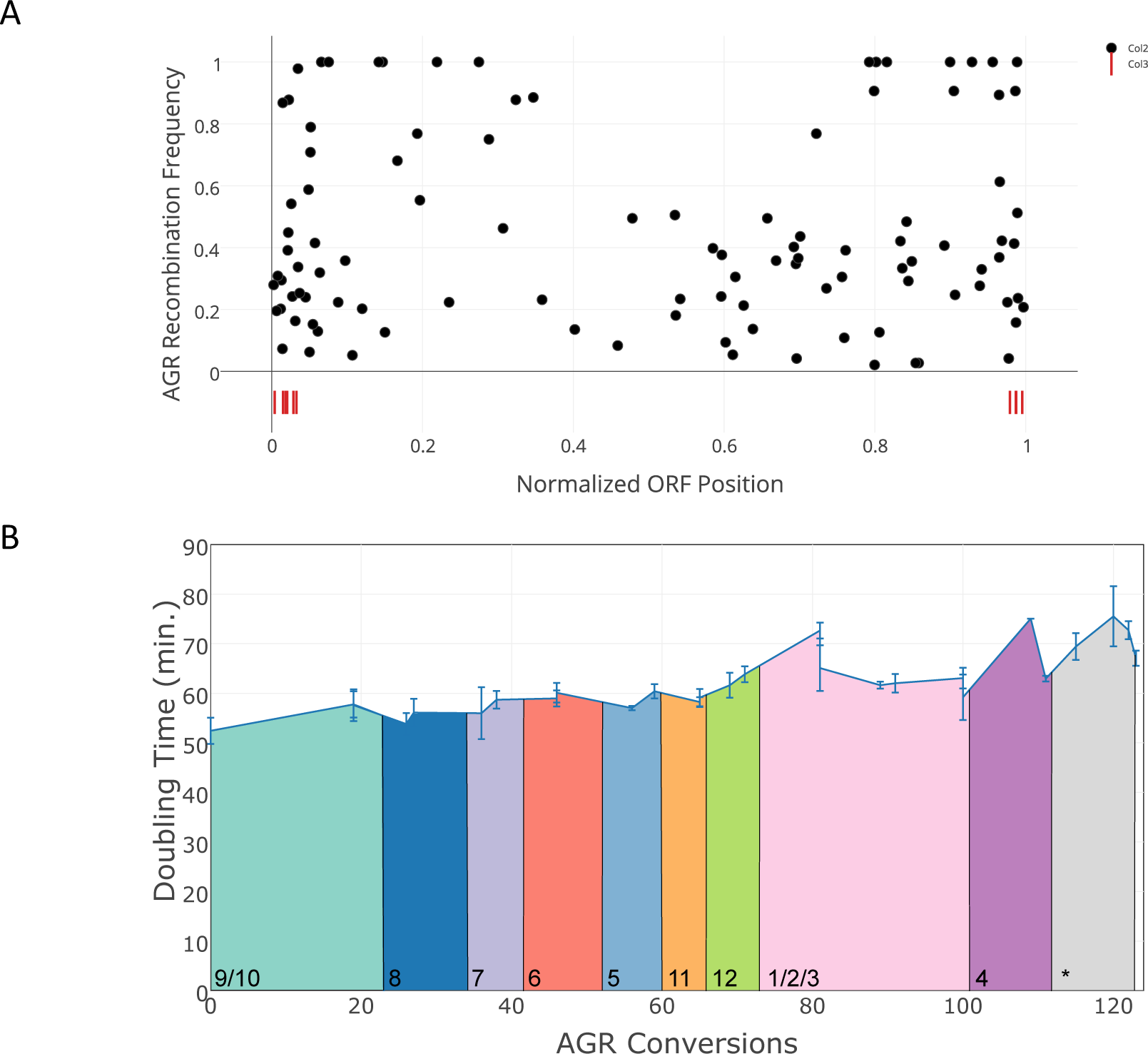
Analysis of attempted AGR→CGU replacements. **(A)** AGR recombination frequency (MASC-PCR, n=96 clones per cell population) was plotted versus the normalized ORF position (residue number of the AGR codon divided by the total length of the ORF). Failed AGR→CGU conversions are indicated using vertical red lines below the x-axis. **(B)** Doubling time of strains in the C123 lineage in LBl media at 34 °C was determined in triplicate on a 96-well plate reader. Colored bars indicate which set of codons was under construction when a doubling time was determined (coloring based on Figure 1). Each data points represent different stages of strain construction. Alternative codons were identified for 13 recalcitrant AGR codons in our troubleshooting pipeline, and the optimized replacement sequences were incorporated into the final strain (gray section at right, labeled with a ’*’), and the resulting doubling times were measured. Error bars represent standard error of the mean in doubling time from at least three replicates of each strain.

**Figure 3.**
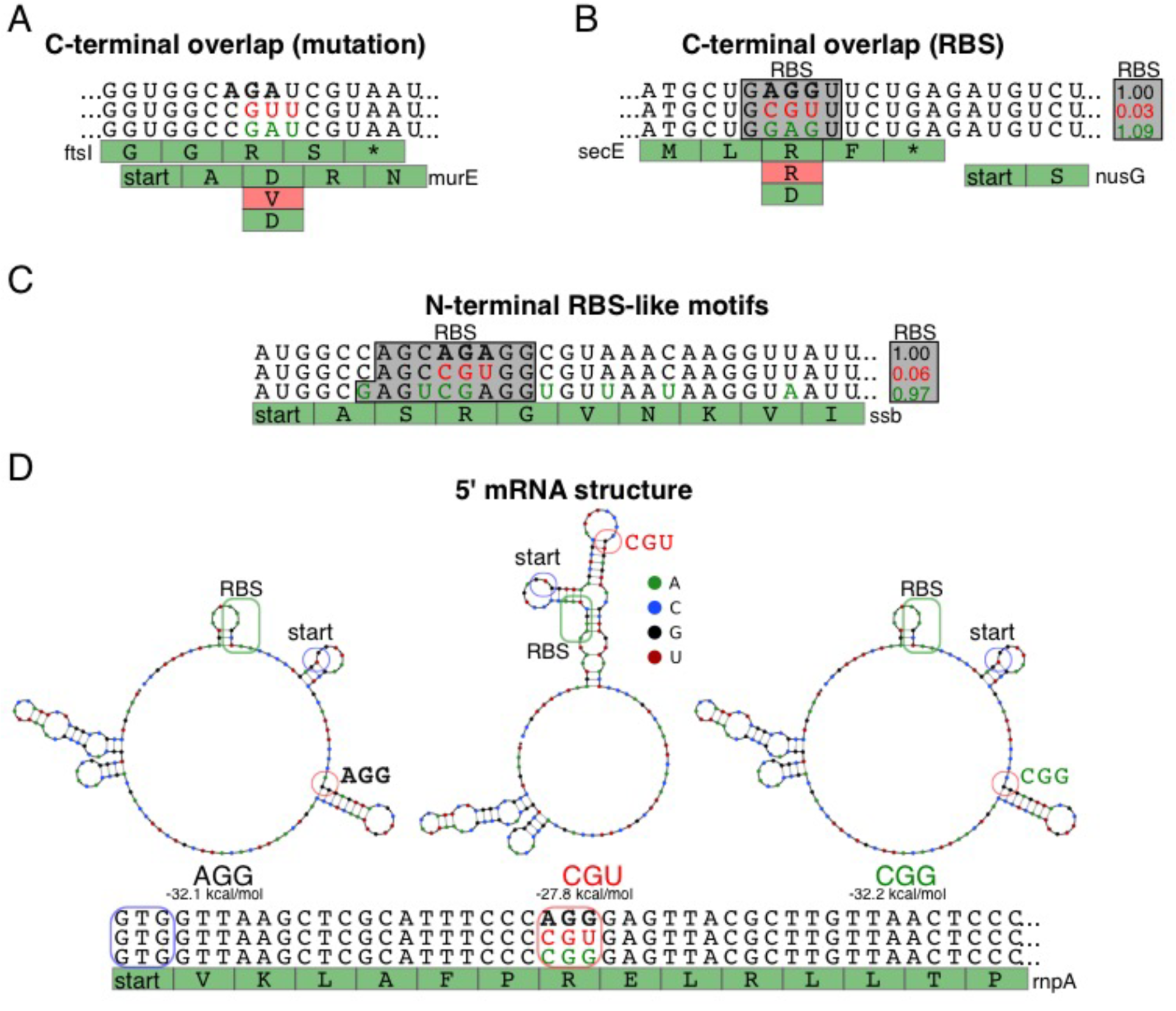
Examples of failure mechanisms for four recalcitrant AGR replacements. Wild type AGR codons are indicated in bold black letters, design flaws are indicated in red letters, and optimized replacement genotypes are indicated in green letters. **(A)** Genes *ftsI* and *murE* overlap with each other. An AGA→CGU mutation in *ftsI* would introduce a non-conservative Asp3Val mutation in *murE*. The amino acid sequence of *murE* was preserved by using an AGA→CGA mutation. **(B)** Gene *secE* overlaps with the RBS for downstream essential gene *nusG*. An AGG→CGU mutation is predicted to diminish the RBS strength by 97% (47). RBS strength is preserved by using a non-synonymous AGG→GAG mutation. **(C)** Gene *ssb* has an internal RBS-like motif shortly after its start codon. An AGG→CGU mutation would diminish the RBS strength by 94%. RBS strength is preserved by using an AGA→CGA mutation combined with additional wobble mutations indicated in green letters. **(D)** Gene *rnpA* has a defined mRNA structure that would be changed by an AGG→CGU mutation. The original RNA structure is preserved by using an AGG→CGG mutation. The RBS (green), start codon (blue) and AGR codon (red) are annotated with like-colored boxes on the predicted RNA secondary structures.

Additionally, the four remaining C-terminal failures included AGR→CGU mutations that disrupt RBS motifs belonging to downstream genes (*secE*_AGG376 for *nusG*, *dnaT*_AGA532 for *dnaC*, and *folC*_AGAAGG1249,1252 for *dedD*, the latter constituting two codonsj. Both *nusG* and *dnaC* are essential, suggesting that replacing AGR with CGU in *secE* and *dnaT* lethally disrupts translation initiation and thus expression of the overlapping *nusG* and *dnaC* (Figure 3B, S2B). Although *dedD* is annotated as non-essential (31), we hypothesized that replacing the AGR with CGU in *folC* disrupted a portion of *dedD* that is essential to the survival of EcM2.1 (*E. coli* K-12). In support of this hypothesis, we were unable to delete the 29 nucleotides of *dedD* that were not deleted by Baba et al. (31) and did not overlap with *folC*, suggesting that this sequence is essential in our strains. The unexpected failure of this conversion highlights the challenge of predicting design flaws even in well-annotated organisms. Consistent with our observation that disrupting these RBS motifs underlies the failed AGR→CGU conversions, we overcame all four design flaws by selecting codons that conserved RBS strength, including a non-synonymous (Arg→Gly) conversion for *secE*.

These lessons, together with previous observations that ribosomes pause during translation when they encounter ribosome binding site motifs in coding DNA sequences (20), provided key insights into the N-terminal AGR→CGU failures. Three of the N-terminal failures (*ssb_AGA10*, *dnaT_AGA10* and *prfB_AGG64*) had RBS-like motifs either disrupted or created by CGU replacement. While *pf_AGG64* is part of the ribosomal binding site motif that triggers an essential frameshift mutation i*nprfB* (21, 38, 39), pausing-motif-mediated regulation of *ssb* and *dnaT* expression has not been reported. Nevertheless, ribosomal pausing data (20) showed that ribosomal occupancy peaks are present directly downstream of the AGR codons for *ssb* and absent for *dnaT* (Figure S3); meanwhile, unsuccessful CGU mutations were predicted to weaken the RBS-like motif for *prfB* and *ssb* and strengthen the RBS-like motif for *dnaT* (Figure 3C, S2C), suggesting a functional relationship between RBS occupancy and cell fitness. Consistent with this hypothesis, successful codon replacements from the troubleshooting pipeline conserve predicted RBS strength compared to the large predicted deviation caused by unsuccessful AGR→CGU mutations (Figure 4, y axis and comparison between orange asterisks and green dots). Interestingly, attempts to replace *dnaT*_AGA10 with either CGN or NNN failed—only by manipulating the wobble position of surrounding codons and conserving the arginine amino acid could *dnaT*_AGA10 be replaced (Figure S2C). These wobble variants appear to compensate for the increased RBS strength caused by the AGA→CGU mutation—RBS motif strength with wobble variants deviated 8-fold from the unmodified sequence, whereas RBS motif strength for AGA→CGU alone deviated 27-fold.

**Figure 4.**
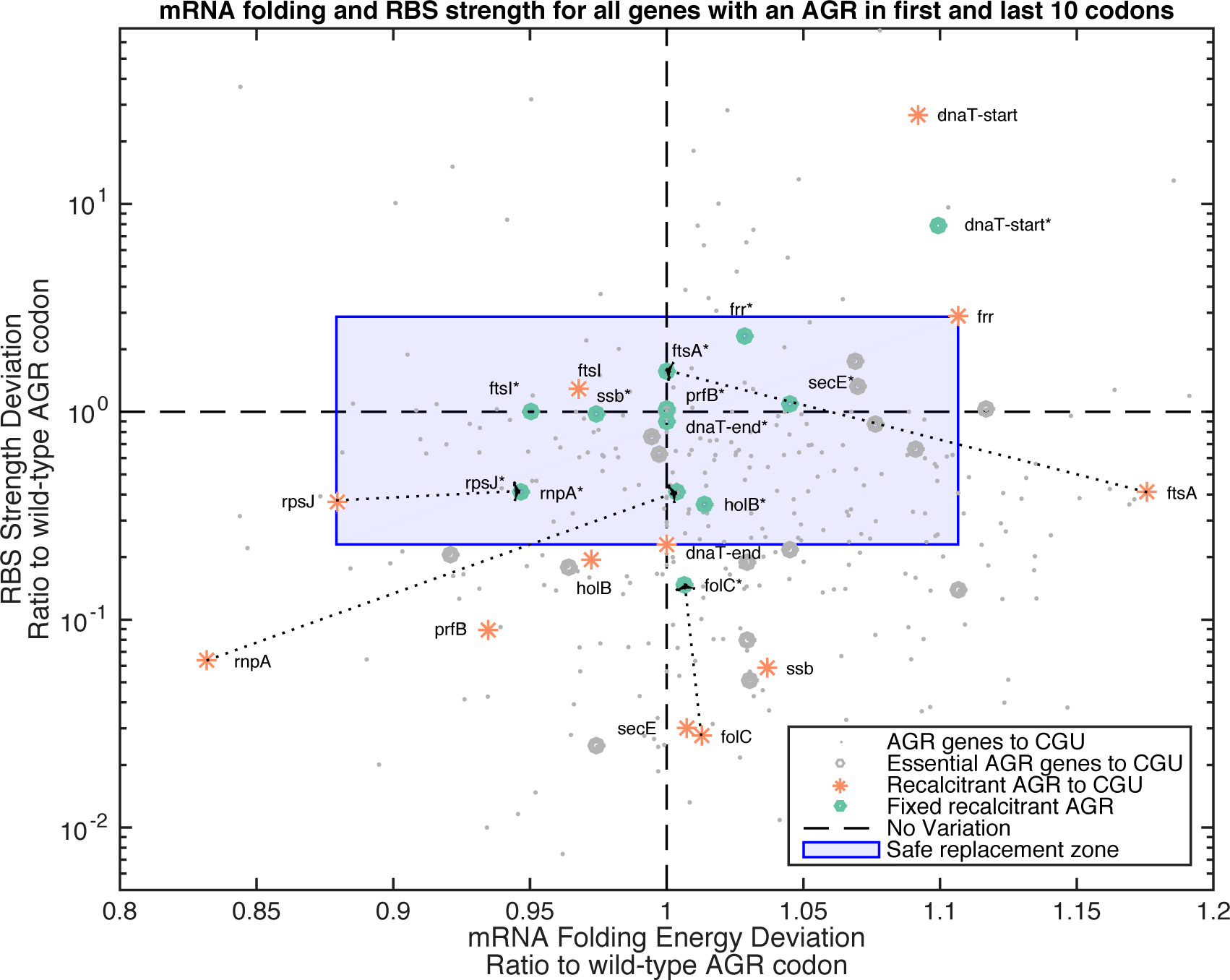
RBS strength and mRNA structure predict synonymous mutation success. Scatter plot showing predicted RBS strength (y-axis, calculated with the Salis ribosome binding site calculator (47)) versus deviations in mRNA folding (x-axis, calculated at 37°C by UNAFold Calculator (41)). Small gray dots represent non-essential genes in *E. coli* MG1655 that have an AGR codon within the first 10 or last 10 codons. Large gray dots represent successful AGR→CGU conversions in the first 10 or last 10 codons of essential genes. Orange asterisks represent unsuccessful AGR→CGU mutations (recalcitrant codons) in essential genes. Green dots represent optimized solutions for these recalcitrant codons. The “safe replacement zone” (blue shaded region) is an empirically defined range of mRNA folding and RBS strength deviations, based on the successful AGR→CGU replacement mutations observed in this study. Most unsuccessful AGR→CGU mutations (Orange asterisks) cause large deviations in RBS strength or mRNA structure that are outside the “safe replacement zone.” Genes *holB* and *ftsI* are two notable exceptions because their initial CGU mutations caused amino acid changes in overlapping essential genes. Gene *folC* corresponds to 2 AGRs. Arrows for four examples of optimized replacement codons (*ftsA, folC, rnpA, rpsJ*) show that deviations in RBS strength and/or mRNA structure are reduced. Arrows are omitted for the remaining 8 optimized replacement codons so as to increase readability.

In order to better understand several remaining N-terminal failure cases that did not exhibit considerable RBS strength deviations (*rnpA_AGG22*, *ftsA_AGA19*, *fTr_AGA16*, and *rpsJ_AGA298*), we examined other potential nucleic acid determinants of protein expression. Based on the observation that mRNA secondary structure near the 5’ end of Open Reading Frames (ORFs) strongly impacts protein expression (12), we found that these four remaining AGR→CGU mutations changed the predicted folding energy and structure of the mRNA near the start codon of target genes (Figure. 3D, S4). Successful codon replacements obtained from degenerate MAGE oligos reduced the disruption of mRNA secondary structure compared to CGU (Figure 4, green dots). For example, *rnpA* has a predicted mRNA loop near its RBS and start codon that relies on base pairing between both guanines of the AGG codon to nearby cytosines (Figure 3D, S5A). Importantly, only AGG22CGG was observed out of all attempted *rnpA* AGG22CGN mutations, and the fact that only CGG preserves this mRNA structure suggests that it is physiologically important (Figure 3D, S5B-C). In support of this, we successfully introduced a *rnpA* AGG22CUG mutation (Arg→Leu) only when we changed the complementary nucleotides in the stem from CC (base pairs with A<u>GG</u>) to CA (base pairs with C<u>UG</u>), thus preserving the natural RNA structure (Figure S5D) while changing both RBS motif strength and amino-acid identity. Our analysis of all four optimized gene sequences showed reduced deviation in computational mRNA folding energy (computed with UNAFold(40)) compared to the unsuccessful CGU mutations (Figure 4, x-axis orange asterisks and green dots). Similarly, predicted mRNA structure (computed with a different mRNA folding software: NUPACK(41)) for these genes was strongly changed by CGU mutations and corrected in our empirically optimized solutions (Figure S4).

Troubleshooting these 13 recalcitrant codons revealed that mutations causing large deviations from natural mRNA folding energy or RBS strength are associated with failed codon substitutions. By calculating these two metrics for all attempted AGR→CGU mutations, we empirically defined a safe replacement zone (SRZ) inside which most CGU mutations were tolerated (Figure 4, shaded area). The SRZ is defined as the largest multi-dimensional space that contains none of the mRNA folding energy or RBS strength associated AGR→CGU failures (Figure 4, red asterisks). It comprises deviations in mRNA folding energy of less than 10% with respect to the natural codon and deviations in RBS-like motif scores of less than a half log with respect to the natural codon, providing a quantitative guideline for codon substitution. Notably, the optimized solution used to replace the 13 recalcitrant codons always exhibited reduced deviation for at least one of these two parameters compared to the deviation seen with a CGU mutation. Furthermore, solutions to the 13 recalcitrant codons overlapped almost entirely with the empirically-defined SRZ. These results suggest that computational predictions of mRNA folding energy and RBS strength can be used as a first approximation to predict whether a designed mutation is likely to be viable. Developing *in silico* heuristics to predict problematic alleles streamlines the use of *in vivo* genome engineering methods such as MAGE to empirically identify viable replacement codons. Therefore, these heuristics reduce the search space required to redesign viable genomes, raising the prospect of creating radically altered genomes exhibiting expanded biological functions.

Once we had identified viable replacement sequences for all 13 recalcitrant codons, we combined the successful 110 CGU conversions with the 13 optimized codon substitutions to produce strain C123, which has all 123 AGR codons removed from all of its annotated essential genes. C123 was then sequenced to confirm AGR removal and analyzed using Millstone, a publicly available genome resequencing analysis pipeline (42). Two spontaneous AAG (Lys) to AGG (Arg) mutations were observed in the essential genes *pssA* and *cca*. While attempts to revert these mutations to AAG were unsuccessful—perhaps suggesting functional compensation—we were able to replace them with CCG (Pro) in *pssA* and CAG (Gln) in *cca* using degenerate MAGE oligos. The resulting strain, C123a, is the first strain completely devoid of AGR codons in its annotated essential genes (Sequences available online). Although some AGR codons in non-essential genes could unexpectedly prove to be difficult to change, our success at replacing all 123 instances of AGR codons in essential genes provides strong evidence that the remaining 4105 AGR codons can be completely removed from the *E. coli* genome, permitting the unambiguous reassignment of AGR translation function (23).

Kinetic growth analysis showed that the doubling time increased from 52.4 (+/− 2.6) minutes in EcM2.1 (0 AGR codons changed) to 67 (+/− 1.5) minutes in C123a (123 AGR codons changed in essential genes) in lysogeny broth (LB) at 34 °C in a 96-well plate reader (See Materials and Methods). Notably, fitness varied significantly during C123 strain construction (Figure 2B). This may be attributed to codon deoptimization (AGR→CGU) and compensatory spontaneous mutations to alleviate fitness defects in a mismatch repair deficient (*mutS-*) background. Overall the reduced fitness of C123a may be caused by on-target (AGR→CGU) or off-target (spontaneous mutations) that occurred during strain construction. In this way, *mutS* inactivation is simultaneously a useful evolutionary tool and a liability. Final genome sequence analysis revealed that along with the 123 desired AGR conversions, C123a had 419 spontaneous non-synonymous mutations not found in the EcM2.1 parental strain (Figure S10). Of particular interest was the mutation *argU_G15A*, located in the D arm of tRNA^Arg^ (*argU*), which arose during CoS-MAGE with AGR set 4. We hypothesized that *argU_G15A* compensates for increased CGU demand and decreased AGR demand, but we observed no direct fitness cost associated with reverting this mutation in C123, and *argU_G15A* does not impact aminoacylation efficiency *in vitro* or aminoacyl-tRNA pools in vivo (Figure S6). Consistent with Mukai et al. and Baba et al. (25, 31), argW (tRNA^Arg^ ccu; decodes AGG only) was dispensable in C123a because it can be complemented by argU (tRNA^Arg^UCU; decodes both AGG and AGA). However, argU is the only *E. coli* tRNA that can decode AGA and remains essential in C123a probably because it is required to translate the AGR codons for the rest of the proteome (23).

To evaluate the genetic stability of C123a after removal of all AGR codons from all the known essential genes, we passaged C123a for 78 days (640 generations) to test whether AGR codons would recur and/or whether spontaneous mutations would improve fitness. After 78 days, no additional AGR codons were detected in a sequenced population (sequencing data available at https://github.com/churchlab/agr_recoding) and doubling time of isolated clones ranged from 22% faster to 22% slower than C123a (n=60).

To gain more insight into how local RBS strength and mRNA folding impact codon choice, we performed an evolution experiment to examine the competitive fitness of all 64 possible codon substitutions at each of AGR codons (Table S2). While MAGE is a powerful method to explore viable genomic modifications *in vivo*, we were interested in mapping the fitness cost associated with less-optimal codon choices, requiring codon randomization depleted of the parental genotype, which we hypothesized to be at or near the global fitness maximum. To do this, we developed a method called CRAM (<u>Cr</u>ispr-<u>A</u>ssisted-<u>M</u>AGE). First, we designed oligos that changed not only the target AGR codon to NNN, but also made several synonymous changes at least 50 nt downstream that would disrupt a 20 bp CRISPR target locus. MAGE was used to replace each AGR with NNN in parallel, and CRISPR/cas9 was used to deplete the population of cells with the parental genotype. This approach allowed exhaustive exploration of the codon space, including the original codon, but without the preponderance of the parental genotype. Following CRAM, the population was passaged 1:100 every 24 hours for six days, and sampled prior to each passage using Illumina sequencing (see Table S2 & Figure 5).

**Figure 5.**
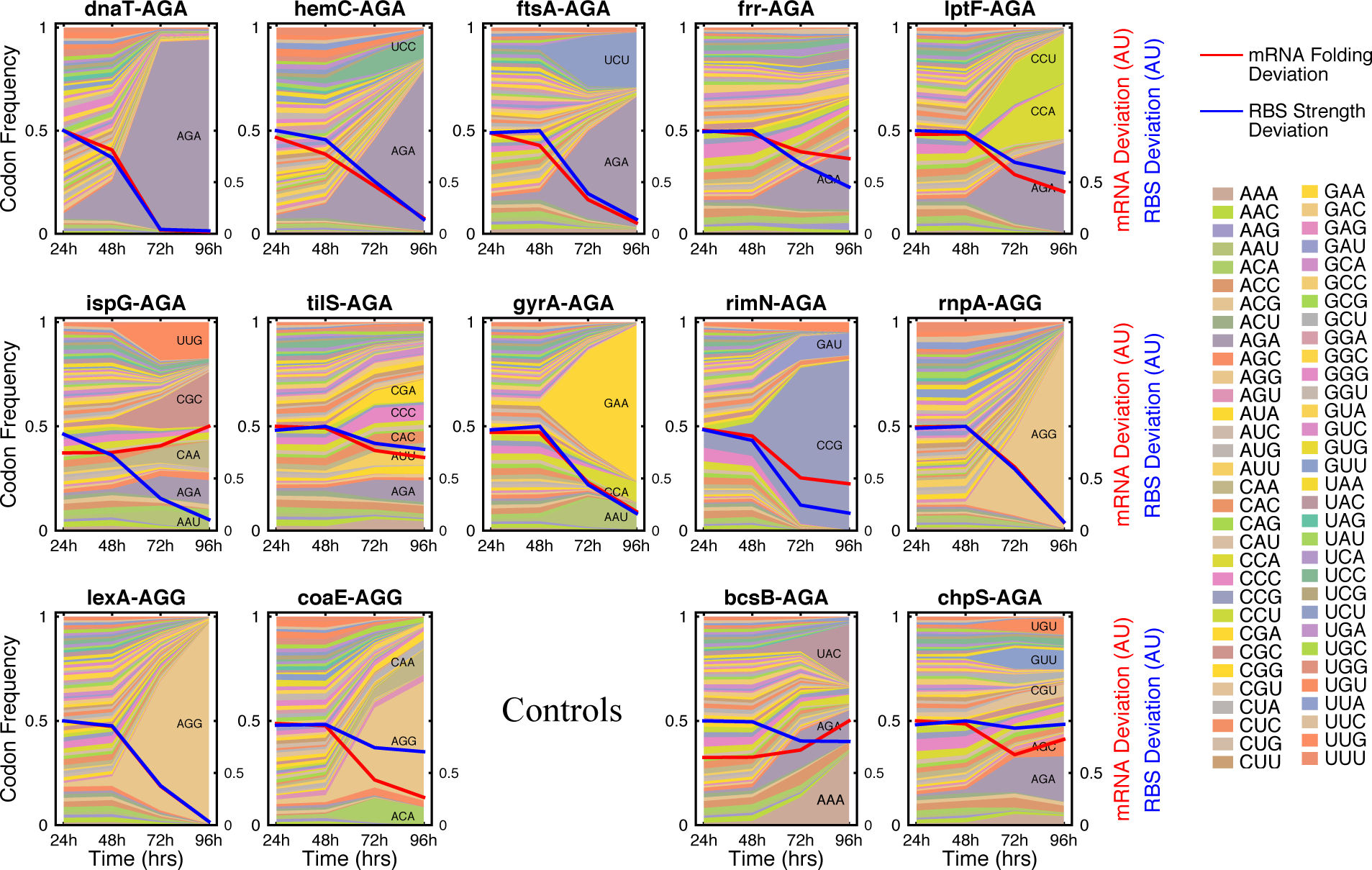
Codon preference of 14 N-terminal AGR codons. CRAM (Crispr-Assisted MAGE) was used to explore codon preference for several AGR codons located within the first 10 codons of their CDS. Briefly, MAGE was used to diversify a population by randomizing the AGR of interest, then CRISPR/Cas9 was used to deplete the parental (unmodified) population, allowing exhaustive exploration of all 64 codons at a position of interest. Thereafter codon abundance was monitored over time by serially passaging the population of cells and sequencing using an Illumina MiSeq. The left y-axis (Codon Frequency) indicates relative abundance of a particular codon (stacked area plot)). The right y-axis indicates the combined deviations in mRNA folding structure (red line) and internal RBS strength (blue line) in arbitrary units (AU) normalized to 0.5 at the initial timepoint. 0 means no deviation from wild type. The horizontal axis indicates the experimental time point in hours at which a particular reading of the population diversity was obtained. Genes *bcsB* and *chpS* are non-essential in our strains and thus serve as controls for AGR codons that are not under essential gene pressure.

Sequencing 24 hours after CRAM showed that all codons were present (including stop codons) (Figure S7), validating the method as a technique to generate massive diversity in a population. All sequences for further analysis were amplified by PCR with allele-specific primers containing the changed downstream sequence. Subsequent passaging of these populations revealed many gene-specific trends (Figure 5, S7, S8). Notably, all codons that required troubleshooting (*dnaT*_AGA10, *ftsA*_AGA19, *frr*_AGA16, *rnpA*_AGG22) converged to their wild-type AGR codon, suggesting that the original codon was globally optimized. For all cases in which an alternate codon replaced the original AGR, we computed the predicted deviation in mRNA folding energy and local RBS strength (as a proxy for ribosome pausing) for these alternative codons and compared these metrics to the evolution of codon distribution at this position over time. We also computed the fraction of sequences that fall within the SRZ inferred from Figure 4 (see Methods). CRAM initially introduced a large diversity of mRNA folding energies and RBS strengths, but these genotypes rapidly converged toward parameters that are similar to the parental AGR values in many cases (Figure 5, overlays). Codons that strongly disrupted predicted mRNA folding and internal RBS strength near the start of genes were disfavored after several days of growth, suggesting that these metrics can be used to predict optimal codon substitutions *in silico*. In contrast, non-essential control genes *bcsB* and *chpS* did not converge toward codons that conserved RNA structure or RBS strength, supporting the conclusion that the observed conservation in RNA secondary structure and RBS strength is biologically relevant for essential genes. Interestingly, *tilS_AGA19* was less sensitive to this effect, suggesting that codon choice at that particular position is not under selection. Additionally, the average internal RBS strength for the *ispG* populations converged towards the parental AGR values whereas mRNA folding energy averages did not, suggesting that this position in the gene may be more sensitive to RBS disruption rather than mRNA folding. Gene *lptF* followed the opposite trend.

Interestingly, several genes (*lptF*, *ispG*, *tilS*, *gyrA* and *rimN*) preferred codons that changed the amino acid identity from Arg to Pro, Lys, or Glu, suggesting that non-coding functions trump amino acid identity at these positions. Importantly, all successful codon substitutions in essential genes fell within the SRZ (Figure 6), validating our heuristics based on an unbiased test of all 64 codons. Meanwhile non-essential control gene *chpS* exhibited less dependence on the SRZ. Based on these observations, while global codon bias may be affected by tRNA availability (6, 43-45), codon choice at a given position may be defined by at least 3 parameters: (1) amino acid sequence, (2) mRNA structure near the start codon and RBS (3) RBS-mediated pausing. In some cases, a subset of these parameters may not be under selection, resulting in an evolved sequence that only converges for a subset of the metrics. In other cases, all metrics may be important, but the primary nucleic acid sequence might not have the flexibility to accommodate all of them equally, resulting in codon substitutions that impair cellular fitness.

**Figure 6.**
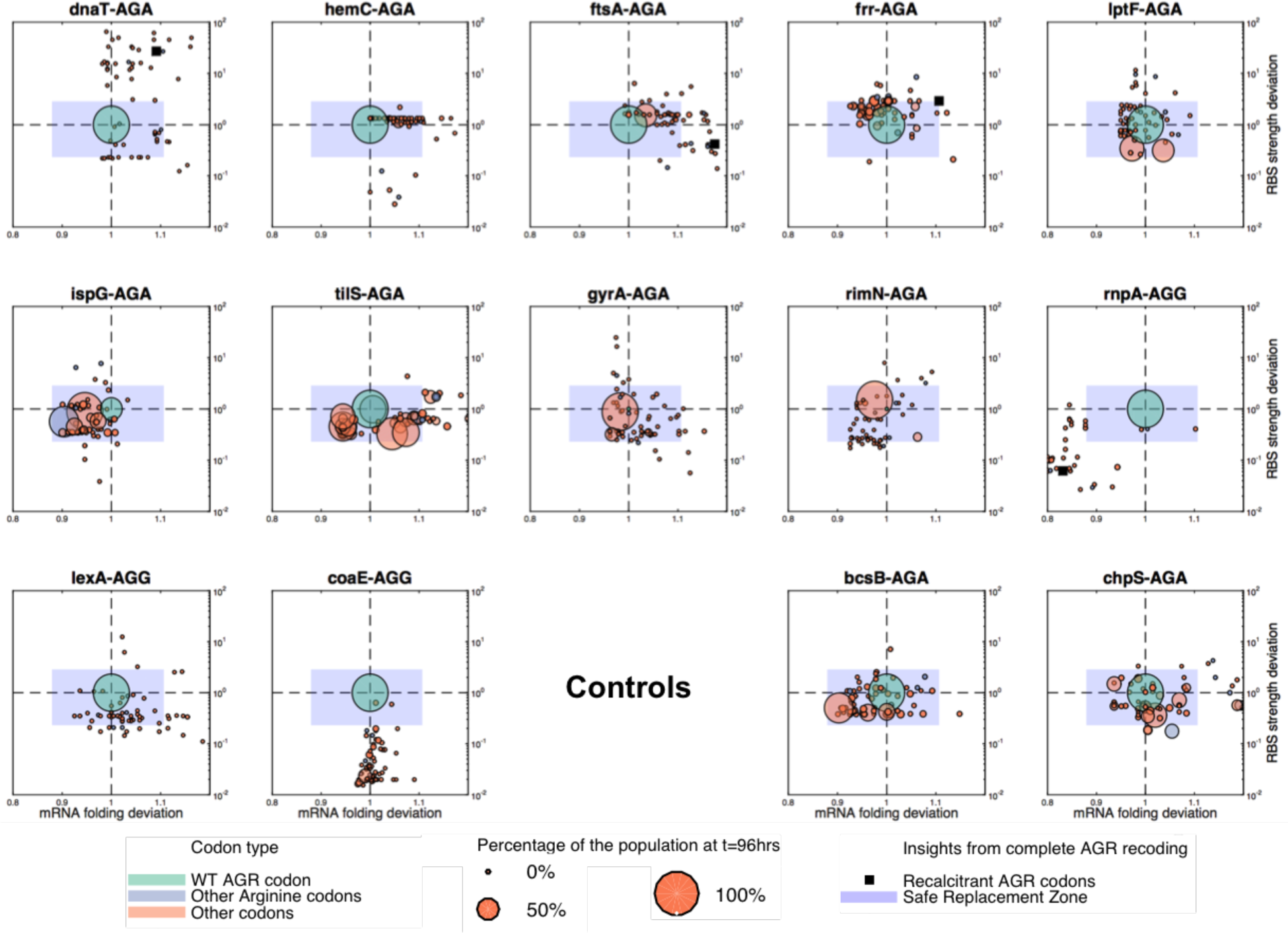
RBS strength and mRNA structure predict codon preference of 14 N-terminal codon substitutions. Scatter plot showing the results of the CRAM experiment (Figure 5). Each panel represents a different gene. The Y-axis represents RBS strength deviation (calculated with the Salis ribosome binding site calculator (47)) while the X-axis shows deviations in mRNA folding energy (x-axis, calculated at 37°C by UNAFold Calculator (41)). Codon abundance at the intermediate time point (t=72hrs, chosen to show maximal diversity after selection) is represented by the dot size. Green dots represent the WT codon. Blue dots represent synonymous AGR codons. Orange dots represent the remaining 58 non-synonymous codons, which may introduce non-viable amino acid substitutions. Black squares represent unsuccessful AGR→CGU conversions observed in the genome-wide recoding effort (Table 1, Figure 1). The “safe replacement zone” (blue shaded region) is the empirically defined range of mRNA folding and RBS strength deviations, based on the successful AGR→CGU replacement mutations observed in this study (Figure 3). Genes *bcsB* and *chpS* are non-essential in our strains and thus serve as controls for AGR codons that are not under essential gene pressure.

These rules were used to generate a draft genome *in silico* with all AGR codons replaced genome-wide, reducing by almost fourfold the number of predicted design flaws (e.g., synonymous codons with metrics outside of the SRZ) compared to the naïve replacement strategy (Figure 7, Figure S9, Table S7, see Methods). Furthermore, predicting recalcitrant codons provides hypotheses that can be rapidly tested *in vivo* using MAGE. Successful replacement sequences can then be implemented together in a redesigned genome. Encouragingly, since all newly predicted design flaws occur in non-essential genes, they would be less likely to impact fitness unless (1) despite the “non-essential” annotation, the gene is actually essential or quasi-essential (i.e., inactivation would impair growth), or (2) the codon in a non-essential gene impacts the expression of a neighboring essential gene (e.g., impacts an RBS motif or RNA structure). While incorrect genome annotations can only be addressed empirically (as demonstrated with gene *dedD*), further analysis reveals that AGR codons in non-essential genes should rarely impact annotated essential genes. In *E. coli* MG1655, only three AGR codons in non-essential genes overlap with the initial mRNA and RBS motifs of essential genes, and at least one synonymous CGN codon is predicted to obey the SRZ for all three cases. Furthermore, even if all synonymous mutations were to disobey the SRZ, since disruption of non-essential gene function should not compromise viability, it is expected that non-synonymous mutations in non-essential genes would be viable as long as they conserve crucial motifs impacting expression of the essential gene. Importantly, we confirmed by MAGE that AGR→CGU codon replacement was possible for 2 of these 3 cases and that an alternative synonymous solution could be found in the remaining case (see Methods).

**Figure 7.**
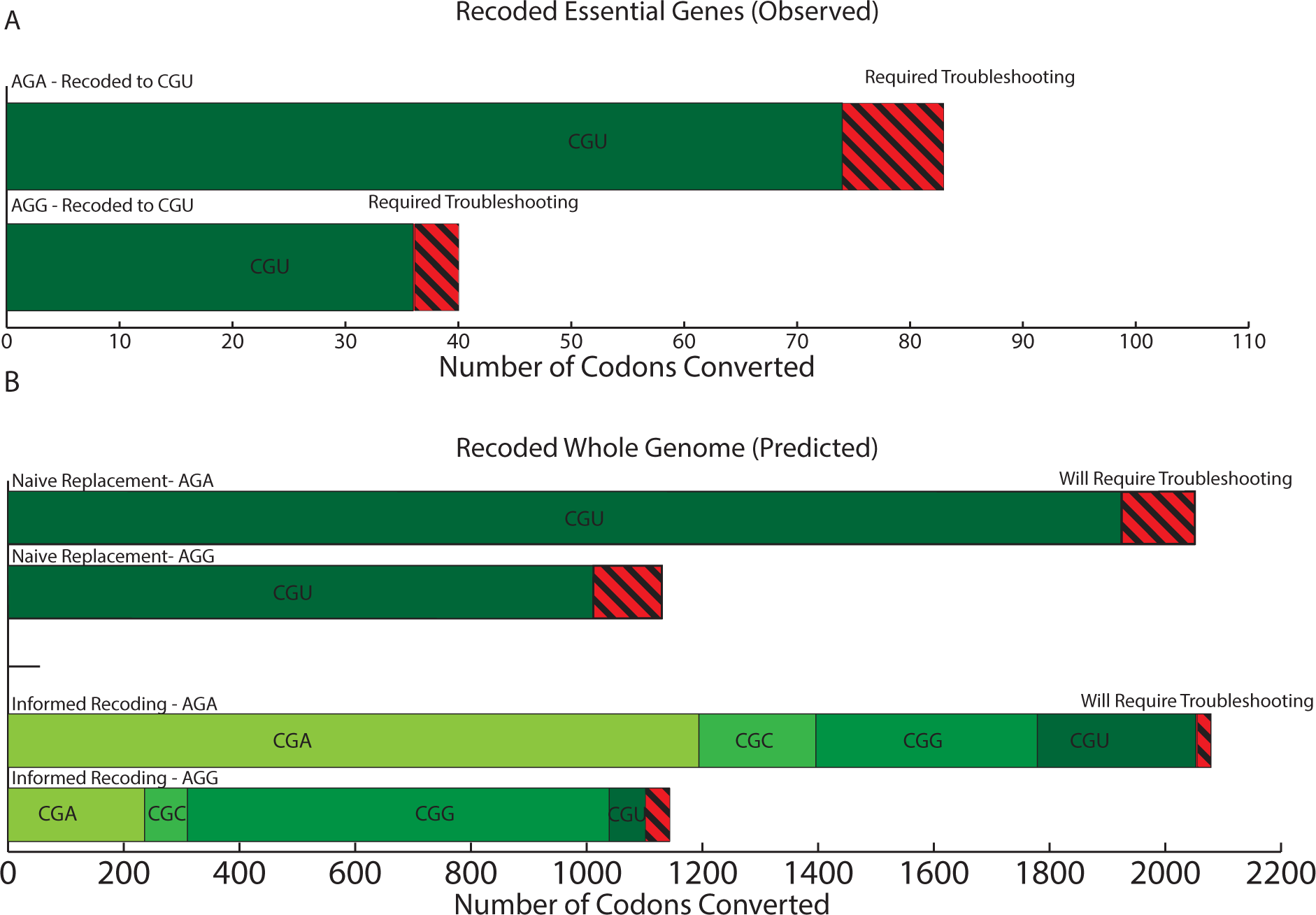
Predicting optimal replacements for AGR codons reduces the number of codons that are predicted to require troubleshooting. **(A)** Empirical data from the construction of C123. 110 AGR codons were successfully recoded to CGU (green), and 13 recalcitrant AGR codons required troubleshooting (red, striped). **(B)** Predicted recalcitrant codons (codons for which no CGN alternatives fall within the SRZ in Figure 4) for replacing all instances of the AGR codons genome-wide. The reference genome used for this analysis had insertion elements and prophages removed (48) to reduce total nucleotides synthesized and to increase genome stability, leaving 3222 AGR codons to be replaced (see Methods). Our analysis predicts that replacing all instances of AGR with CGU would have resulted in 229 failed conversions (‘Naive Replacement’, red striped). However, implementing the rules from this work (‘Informed Replacement’) to identify the best CGN alternative reduces the predicted failure rate from 7.1% (229/3222), to 2.0% (64/3222AGR) of which only a small subset will have a direct impact on fitness due to their location in non-essential genes. In such cases, MAGE with degenerate oligos could be used to empirically identify replacement codons as we have demonstrated herein. Each specific synonymous CGN is identified with a unique shade of green and is labeled inside of its respective section.

To conclude, comprehensively removing all instances of AGR codons from all *E. coli* essential genes revealed 13 design flaws which could be explained by a disruption in coding DNA Sequence, RBS-mediated translation initiation, RBS-mediated translation pausing, or mRNA structure. While the importance of each factor has been reported, our work systematically explores to what extent and at what frequency they impact genome function. Furthermore, our work establishes quantitative guidelines to reduce the chance of designing non-viable genomes. Although additional factors undoubtedly impact genome function, the fact that these guidelines captured all instances of failed synonymous codon replacements (Figure 4) suggests that our genome design guidelines provide a strong first approximation of acceptable modifications to the primary sequence of viable genomes. These design rules coupled with inexpensive DNA synthesis will facilitate the construction of radically redesigned genomes exhibiting useful properties such as biocontainment, virus resistance, and expanded amino acid repertoires (46).

## Acknowledgments

This work was supported by the US Department of Energy [DE-FG02-02ER63445]; by the US Defense Advanced Research Projects Agency [N66001-12-C-4211 to F.J.I. and D.S.]; by the National Institute of General Medical Sciences [GM22854-42 to D.S.]; by a US Department of Defense NDSEG Fellowship (to M.J.L. and G.K.); by a US National Science Foundation Graduate Research Fellowship (to D.B.G.); by the Lynch Foundation (to M.M.L.); by an Amazon AWS in Education Grant Award (to G.K.); and by the Arnold & Mabel Beckman Foundation and DuPont, Inc. (to F.J.I.). Funding for open access charge: US Department of Energy [DE-FG02-02ER63445]. Stephanie Yaung generously provided CRISPR plasmids and technical support. A patent application has been filed by Harvard University relating to aspects of the work described in this manuscript. In the interests of transparency, we wish to mention that G.M.C. is a founder of Enevolv Inc. and Gen9bio (neither of which was involved in this study). Other potentially relevant financial interests are listed at http://arep.med.harvard.edu/gmc/tech.html.

## Materials and Methods

Supplementary Materials

Figures S1-S10

Tables S1-S5

### Supplementary Materials

This section includes the actual text of the Supplementary Materials, which can include any or all of the preceding items, and figure captions and tables that can easily be incorporated into one supplementary material file. Please edit the list above as appropriate and include it at the end of your main paper. If there are additional files that cannot be easily accommodates (e.g., movies or large tables), please include captions here.

#### Materials and Methods

##### Strains & Culture Methods

The strains used in this work were derived from EcM2.1 (*Escherichia coli* MG1655 *mutS_mut dnaG_Q576A exoX_mut xonA_mut xseA_mut 1255700::tolQRA* Δ(*ybhB-bioAB*)::[λcI857 N(*cro-ea59*)::*tetR-bla*]) (33). Liquid culture medium consisted of the Lennox formulation of Lysogeny broth (LB^L−^; 1% w/v bacto tryptone, 0.5% w/v yeast extract, 0.5% w/v sodium chloride) (49) with appropriate selective agents: carbenicillin (50 μg/mL) and SDS (0.005% w/v). For *tolC* counter-selections, colicin E1 (colE1) was used at a 1:100 dilution from an in-house purification (50) that measured 14.4 μg_protein_/μL (22, 36), and vancomycin was used at 64 μg/mL. Solid culture medium consisted of LB^L^ autoclaved with 1.5% w/v Bacto Agar (Fisher), containing the same concentrations of antibiotics as necessary. ColE1 agar plates were generated as described previously (33). Doubling times were determined on a Biotek Eon Microplate reader with orbital shaking at 365 cpm at 34 °C overnight, and analyzed using a matlab script available on GitHub (https://github.com/churchlab/agr_recoding).

##### Oligonucleotides, Polymerase Chain Reaction, and Isothermal Assembly

A complete table of MAGE oligonucleotides and PCR primers can be found in Table S1.

PCR products used in recombination or for Sanger sequencing were amplified with Kapa 2G Fast polymerase according to manufacturer’s standard protocols. Multiplex allele-specific PCR (mascPCR) was used for multiplexed genotyping of AGR replacement events using the KAPA2G Fast Multiplex PCR Kit, according to previous methods (22, 51). Sanger sequencing reactions were carried out through a third party (Genewiz). CRAM plasmids were assembled from plasmid backbones linearized using PCR (52), and CRISPR/PAM sequences obtained in Gblocks from IDT, using isothermal assembly at 50 °C for 60 minutes. (53).

##### Lambda Red Recombinations, MAGE, & CoS-MAGE

λ Red recombineering, MAGE, and CoS-MAGE were carried out as described previously (33, 54). In singleplex recombinations, the MAGE oligo was used at 1 μM, whereas the co-selection oligo was 0.2 μM and the total oligo pool was 5 μM in multiplex recombinations (7-14 oligos).. When double-stranded PCR products were recombined (e.g., *tolC* insertion), 100 ng of double-stranded PCR product was used. Since we used CoS-MAGE with *tolC* selection to replace target AGR codons, each recombination was paired with a control recombined with water only to monitor *tolC* selection performance. The standard CoS-MAGE protocol for each oligo set was to insert *tolC*, inactivate *tolC*, reactivate *tolC*, and delete *tolC*. MascPCR screening was performed at the *tolC* insertion, inactivation and deletion steps. All λ Red recombinations were followed by a recovery in 3 mL LBL followed by a SDS selection (*tolC* insertion, *tolC* activation) or ColE1 counter-selection (*tolC* inactivation, *tolC* deletion) that was carried out as previously described (33).

##### General AGR replacement strategy

AGR codons in essential genes were found by cross-referencing essential gene annotation according to two complementary resources (31, 55) to find the shared set (107 coding regions), which contained 123 unique AGR codons (82 AGA, 41 AGG). We used optMAGE (35, 54) to design 90-mer oligos (targeting the lagging strand of the replication fork) that convert each AGR to CGU. We reduced the total number of AGR replacement oligos to 119 by designing oligos to encode multiple edits where possible, maintaining at least 20 bp of homology on the 5’ and 3’ ends of the oligo. The oligos were then pooled based on chromosomal position into twelve MAGE oligo sets of varying complexity (minimum: 7, maximum: 14) such that a single marker (*tolC*) could be inserted at most 564,622 bp upstream relative to replication direction for all targets within a given set. We then identified *tolC* insertion sites for each of the twelve pools either into intergenic regions or non-essential genes that met the distance criteria for a given pool. See Table 1 for descriptors for each of the 12 oligo pools.

##### Troubleshooting strategy

A recalcitrant AGR was defined as one that was not converted to CGU in one of at least 96 clones picked after the third step of the conversion process. The recalcitrant AGR codon was then triaged for troubleshooting (Fig. 3A**) in the parental strain (EcM2.1). First, the sequence context of the codon was examined for design errors or potential issues, such as misannotation or a disrupted RBS for an overlapping gene. In most cases, corrected oligos could be easily designed and tested. If no such obvious redesign was possible, we attempted to replace AGR with CGN mutations. If attempting to replace AGR with CGN failed to give recombinants, we tested compensatory, synonymous mutations in a 3 amino acid window around the recalcitrant AGR. If needed, we finally relaxed synonymous stringency by recombining with oligos encoding AGR-to-NNN mutations.

After each step in the troubleshooting workflow, we screened 96 clones from 2 successive CoS-MAGE recombinations using allele specific PCR with primers that hybridize to the wildtype genotype. Sequences that failed to yield a wild-type amplicon were Sanger sequenced to confirm conversion. We also measured doubling time of all clones in LBL to pair sequencing data with fitness data, and chose the recombined clone with the shortest doubling time. Doubling time was determined by obtaining a growth curve on a Biotek plate reader (either an Eon or H1), and analyzed using web-based open source genome resequencing software available on GitHub at https://github.com/churchlab/millstone. This genotype was then implemented in the complete strain at the end of strain construction using MAGE, and confirmed by MASC-PCR screening.

##### AGR codons in non-essential genes with impact on essential genes

In *E. coli* MG1655, only three AGR codons in non-essential genes overlap with the initial mRNA and RBS motifs of essential genes, and at least one synonymous CGN codon is predicted to obey the SRZ for all three cases. As in the troubleshooting pipeline, we attempted to replace AGR with CGT mutations using MAGE. After 4 cycles of MAGE, cells were plated and 96 clones were screened. Synonymous codon replacement was possible for genes *rffT* and *mraW*, but not for gene *yidD*. We then relaxed synonymous stringency by recombining with oligos encoding AGR-to-NNN mutations for gene *yidD* and found multiple alternative solutions including CGA, UGA, GUG, GCG and TAA. Importantly, the synonymous CGA alternative solutions disrupted less RBS strength and mRNA folding that CGU (see table S7) further confirming our rules as useful guidelines.

##### mRNA folding and RBS strength computations

A custom Python pipeline (available at https://github.com/churchlab/agr_recoding) was used to compute mRNA folding and RBS strength value for each sequence. mRNA folding was based on the UNAFold calculator (40) and RBS strength on the Salis calculator (47). The parameters for mRNA folding are the temperature (37°C) and the window used which was an average between −30:+100nt and −15:+100nt around the start site of the gene and was based on Goodman et al., 2013. The only parameter for RBS strength is the distance between RBS and promoter and we averaged between 9 and 10 nt after the codon of interest based on Li et al., 2012. Data visualization was performed through a custom Matlab code.

For *in silico* predictions on the entire genome, all 3222 AGR in non-phage genes were analyzed using this custom pipeline and data is presented in Table S7. Phage genes were not analyzed to reduce the complexity of the genome, inspired by other reduced genome efforts (48).

##### Whole genome sequencing of strains lacking AGR codons in their essential genes

Sheared genomic DNA was obtained by shearing 130uL of purified genomic DNA in a Covaris E210. Whole genome library prep was carried out as previously described (56). Briefly, 130 uL of purified genomic DNA was sheared overnight in a Covaris E210 with the following protocol: Duty cycle 10%, intensity 5, cycles/burst 200, time 780 seconds/sample. The samples were assayed for shearing on an agarose gel and if the distribution was acceptable (peak distribution ~400 nt) the samples were size-selected by SPRI/Reverse-SPRI purification as described in (56). The fragments were then blunted and p5/p7 adaptors were ligated, followed by fill-in and gap repair (NEB). Each sample was then qPCR quantified using SYBR green and Kapa Hifi. This was used to determine how many cycles to amplify the resulting library for barcoding using P5-sol and P7-sol primers. The resulting individual libraries were quantified by Nanodrop and pooled. The resulting library was quantified by qPCR and an Agilent Tapestation, and run on MiSeq 2x150. Data was analyzed to confirm AGR conversions and to identify off-target mutations using Millstone, an web-based open-source genome resequencing tool.

Seqeunces are available online at https://github.com/churchlab/agr_recoding.

##### NNN-sequencing and CRISPR

CRISPR/Cas9 was used to deplete the wildtype parental genotype by selectively cutting chromosomes at unmodified target sites next to the desired AGR codons changes. Candidate sites were determined using the built-in target site finder in Geneious proximally close to the AGR codon being targeted. Sites were chosen if they were under 50 bp upstream of the AGR codon and could be disrupted with synonymous changes. If multiple sites fulfilled these criteria, the site with the lowest level of sequence similarity to other portions of the genome was chosen. Oligos of a length of ~130 bp were designed for all 14 genes with an AGR codon in the first 30 nt after the translation start site. Those oligos incorporated both an NNN random codon at the AGR position as well as multiple (up to 6) synonymous changes in a CRISPR target site at least 50 nt downstream of an AGR codon. This modifies the AGR locus at the same time as disrupting the CRISPR target site, ensuring randomization of the locus after the parental genotype is deleted.

Specifically, we constructed a plasmid containing the SpCas9 protein gene (Plasmid details (DS-SPcas, Addgene plasmid 48645): cloDF13 origin, specR, proC promoter, SPcas9, unused tracrRNA (with native promoter and terminator), J23100 promoter, 1 repeat (added to facilitate cloning in a spacer onto the same plasmid). We also constructed 14 plasmids containing the guide RNA directed towards the unmodified sequences (Plasmid details (PM-!T4Y): p15a origin, chlor^R^, J23100 promoter, spacer targeting T4, 1 repeat).

For each of 24 genes, five cycles of MAGE were performed with the specific mutagenesis oligo at a concentration of 1uM. CRISPR repeat-spacer plasmids carrying guides designed to target the chosen sites, and were electroporated into each diversified pool after the last recombineering cycle. After 1 hour of recovery, both the SpCas9 and repeat-spacer plasmids were selected for, and passaged in three parallel lineages for each of the 24 AGR codons for 144 hrs. After 2 hours of selection, and at every 24 hour interval, samples were taken and the cells were diluted 1/100 in selective media.

Each randomized population was amplified using PCR primers allowing for specific amplification of strains incorporating the CRISPR-site modifications. The resulting triplicate libraries for each AGR codon were then pooled and barcoded with P5-sol and P7-sol primers, and run on a MiSeq 1x50. Data was analyzed using custom Matlab code, available on https://github.com/churchlab/agr_recoding.

For each gene and each data point, reads were aligned to the reference genome and frequencies of each codon were computed. In figure 5, the mRNA structure deviation (red line) and RBS strength deviation (blue line) in arbitrary units were computed based as the product of the frequencies and the corresponding deviation for each codon.

**Table S1.** Full list and positions, oligo ID, and pool ID (position in the genome)

**Table S2.** Summary of CRAM results

**Table S3.** Kinetic data for argU mutations found in our strain

**Table S4.** Curated summary of mutations found in the final constructed strain, C123a (non-synonymous and high impact mutations only)

**Table S5.** MAGE oligos used for converting 123 AGR codons in essential genes.

**Table S6.** List of recalcitrant AGR→CGU conversions.

**Table S7.** mRNA folding deviation and RBS strength deviation for all AGR codon genome wide.

**Figure S1.**
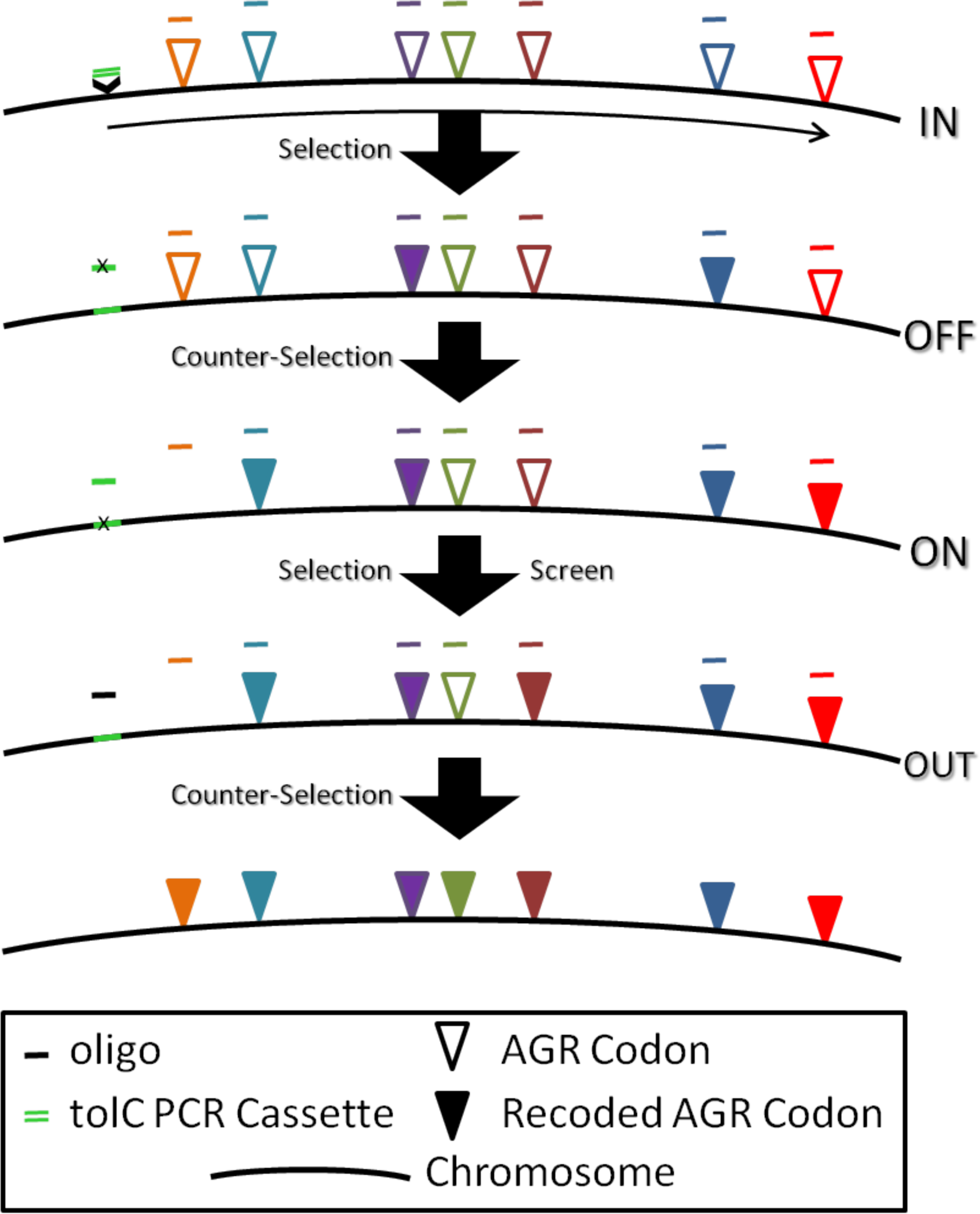
Strategy for replacing each “set” of AGR codons in all of the essential genes of *Escherichia coli (EcM2.1)*. Here the AGR codons are marked with open triangles (various colors). To start, a dual-selectable *tolC* cassette (double green line) is recombined into the genome using lambda red in a multiplexed recombination along with several oligos targeting nearby (<500 kb), downstream AGR loci (various colored lines). Upon selection for *tolC* insertion clones, correctly chosen AGR codons are also observed (filled in triangles) at a higher frequency due to strong linkage between recombination events at *tolC* and other nearby (< 500 kb), downstream AGR loci. Next, a second recombination is carried out using the same AGR conversion oligo pool, but now paired with another oligo to disrupt the *tolC* ORF with a premature stop, after which the *tolC* counter-selection is applied, again enriching the population for AGR conversions. A third, multiplexed recombination then fixes the *tolC* ORF, again targeting AGR loci. After applying the *tolC* selection clones are assayed by MASC-PCR. Assuming most conversions in a given set had been made, the selectable marker would then be removed using a repair oligo in a singleplexed or multiplexed recombination (depending on need). The *tolC* counter-selection is then leveraged to both leave a scarless chromosome and free up the *tolC* cassette for use elsewhere in the genome.

**Figure S2.**
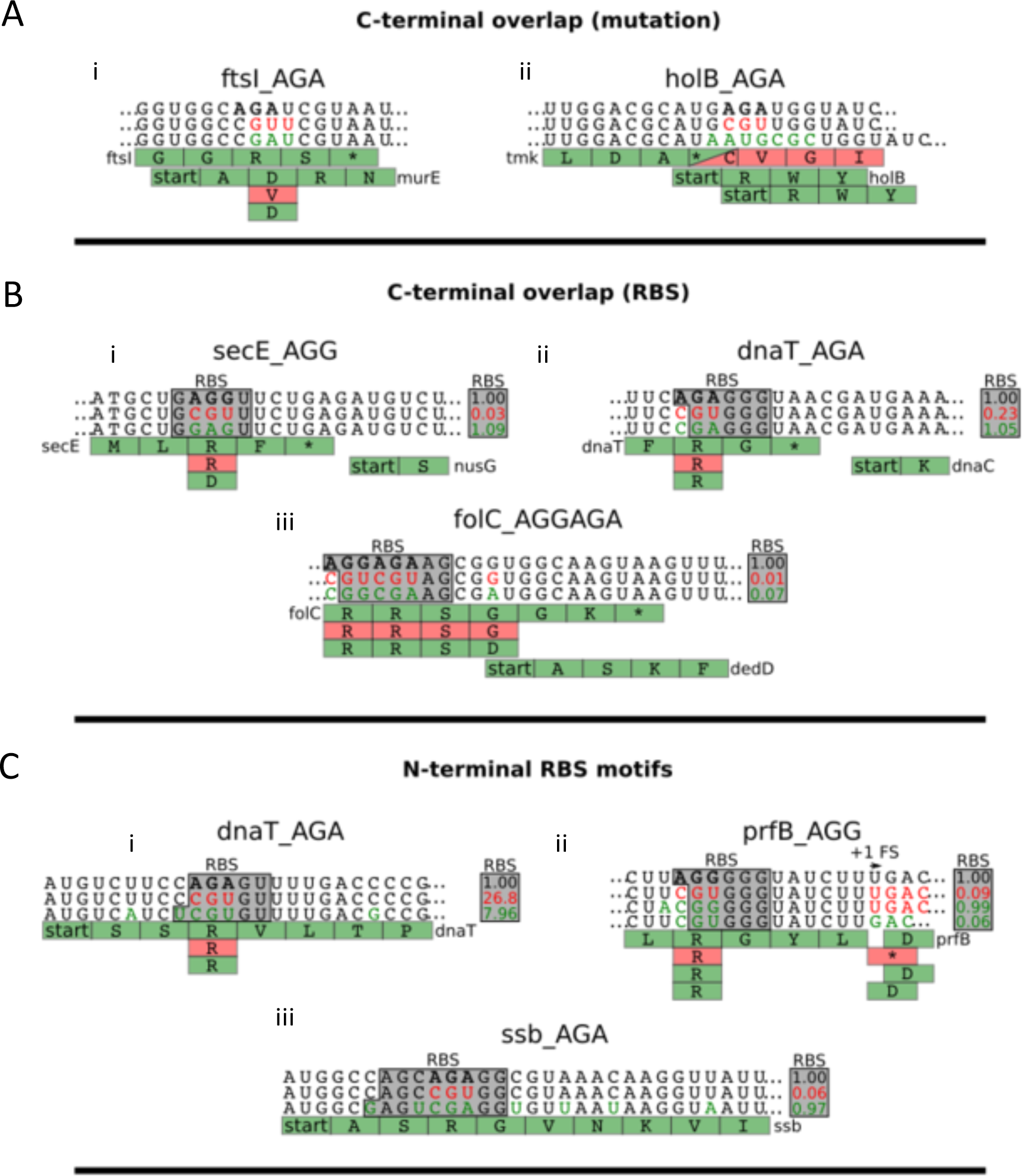
Schematic of 3 different failures cases for recalcitrant AGR→CGU mutations. For each case, the top row is the initial sequence, the middle row is the AGR→CGU mutation and the third row of primary DNA sequence is the optimized solution converged on in troubleshooting. Green boxes below the DNA sequence indicates amino acid sequence in the same order (top is initial, middle results from AGR→CGU, bottom results from troubleshot solution). **A.** C-terminal overlap cases of AGR’s at ends of essential genes with downstream ORF’s. (i) Genes *ftsI* and *murE* overlap with each other. An AGA→CGU mutation in *ftsI* would introduce a non-conservative Asp3Val mutation in *murE*. The amino acid sequence of *murE* was preserved by using an AGA→CGA mutation. (ii) Genes *holB* and *tmk* overlap with each other. An AGA→CGU mutation in *holB* would introduce a non-conservative Stop214Cys mutation in *tmk*. The amino acid sequence of *tmk* was preserved by using an AGA→CGC mutation and adding 3 nucleotides. **B.** C-terminal overlap cases of AGR’s at ends of essential genes with the RBS of a downstream gene. (i) Gene *secE* overlaps with the RBS for downstream essential gene *nusG*. An AGG→CGU mutation would diminish the RBS strength by 97% (47). RBS strength is preserved by using an AGG→AG mutation. (ii) Gene *dnaT* overlaps with the RBS for downstream essential gene *dnaC*. An AGG→CGU mutation would diminish the RBS strength by 77% (47). RBS strength is preserved by using an AGG→CGA mutation. (ii) Gene *folC* overlaps with the RBS for downstream gene *dedD*, shown to be essential in our strain. An AGGAGA→CGUCGU mutation would diminish the RBS strength by 99% (47). RBS strength is preserved by using an AGG→CGGCGA mutation. **C.** N-terminal RBS-like motifs causing recalcitrant AGR conversions at the beginning of essential genes. (i) Gene *dnaT* has an internal RBS-like motif. An AGG→CGU mutation would increase the RBS strength 26 times (47). RBS strength is better preserved by using an AGA→CGU mutation combined with additional wobble mutations. (ii) Gene *prfB* has an internal RBS-like motif. This RBS-like motif is involved in a downstream planned frameshift in prfB (39). Only by removing the frameshift was AGG→CGU mutation possible (leaving a poor RBS-like site). To maintain the frameshift, AGG→CGG mutation and additional wobble was required. In that case, local RBS strength was maintained (fourth row). (iii) Gene *ssb* has an internal RBS-like motif. An AGG→CGU mutation would diminish the RBS strength by 94%. RBS strength is preserved by using an AGA→CGA mutation combined with additional wobble mutations.

**Figure S3.**
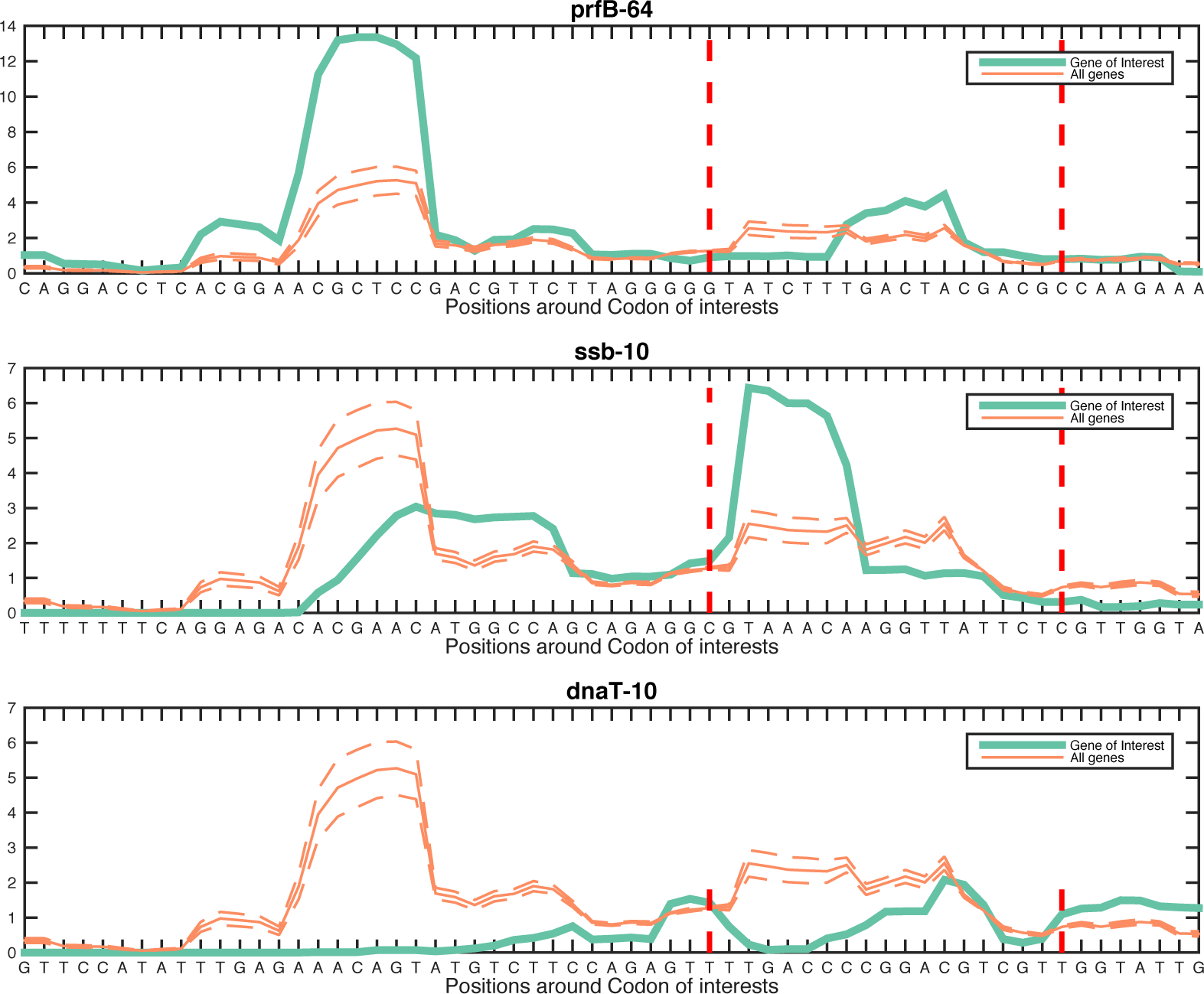
Ribosomal pausing data drawn from previous work (Li et al., 2012) for genes *ssb*, *dnaT* and *prfB*. Green line represent ribosome profiling data for each gene. Orange line is the average for all genes with an AGR codon within the first 30 nucleotides of the annotated start codon. Region between the two vertical red lines indicates zones of interest (centered 12bp after the AGR codon). Interestingly, *prfB* and *ssb* show a peak after the AGR codon, where no peak is observed for *dnaT*. Based on predictions from the Salis calculator (47), replacing AGR with CGU in those 3 cases is believed to disrupt ribosomal pausing (*prfB* and *ssb*) or to introduce ribosomal pausing (*dnaT*).

**Figure S4.**
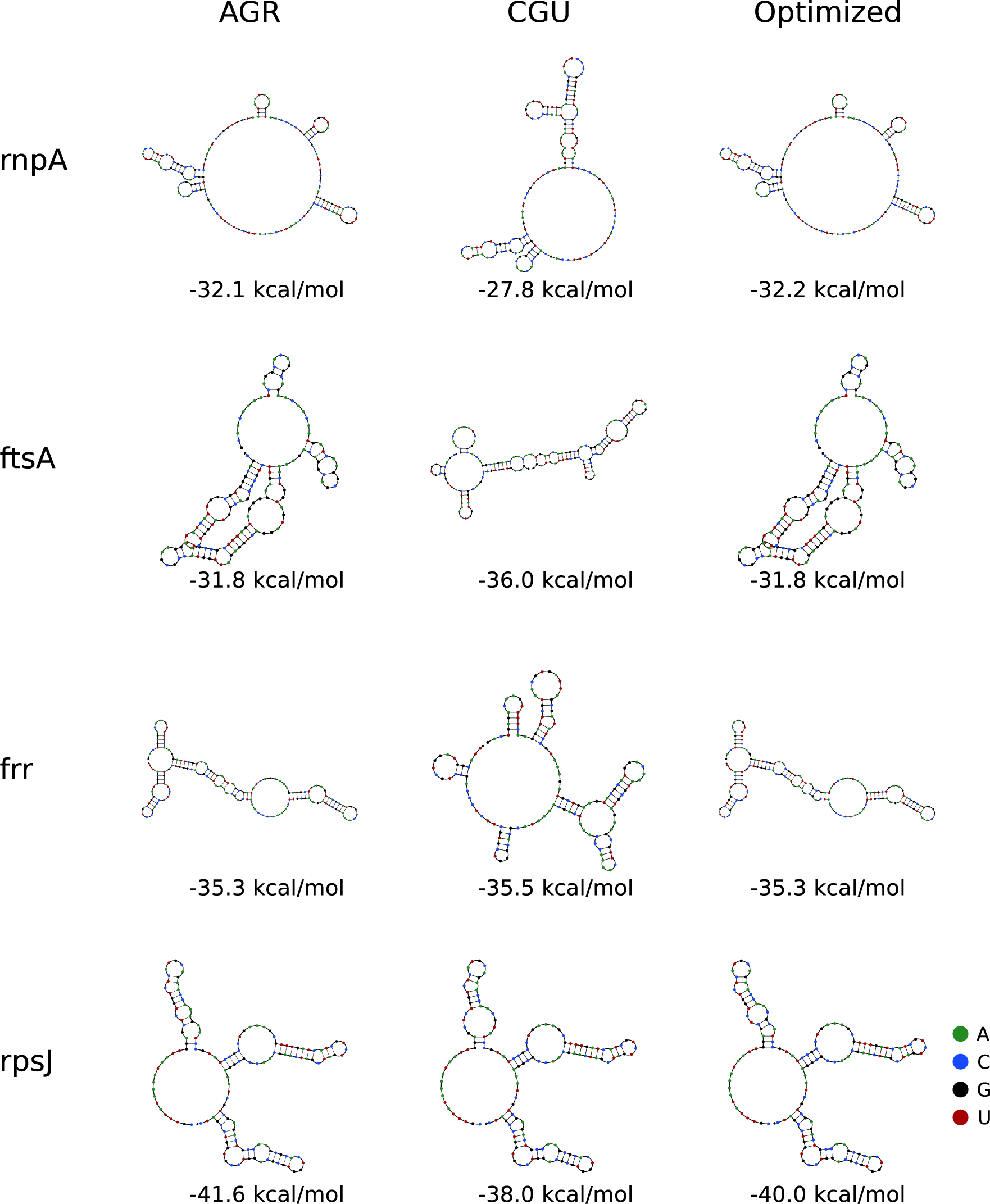
mRNA folding predictions for the 4 recalcitrant AGR→CGU mutations explained by mRNA folding variations. mRNA folding prediction of 100 nucleotides upstream and 30 nt downstream of the start codon using UNAfold (40). Both the shape of the mRNA folding and the folding energy value have to be taken into account to understand failure of the AGR→CGU conversion. ‘AGR’ depicts the predicted, wild-type mRNA, ‘CGU’ is the mRNA folding prediction with an AGR→CGU mutation (generally not observed) and ‘Optimized’ correspond to the mRNA folding prediction of the AGR replacement solution found after *in vivo* troubleshooting. Under each structure, the predicted free energy of folding of the visualized structure is listed in kcal/mol.

**Figure S5.**
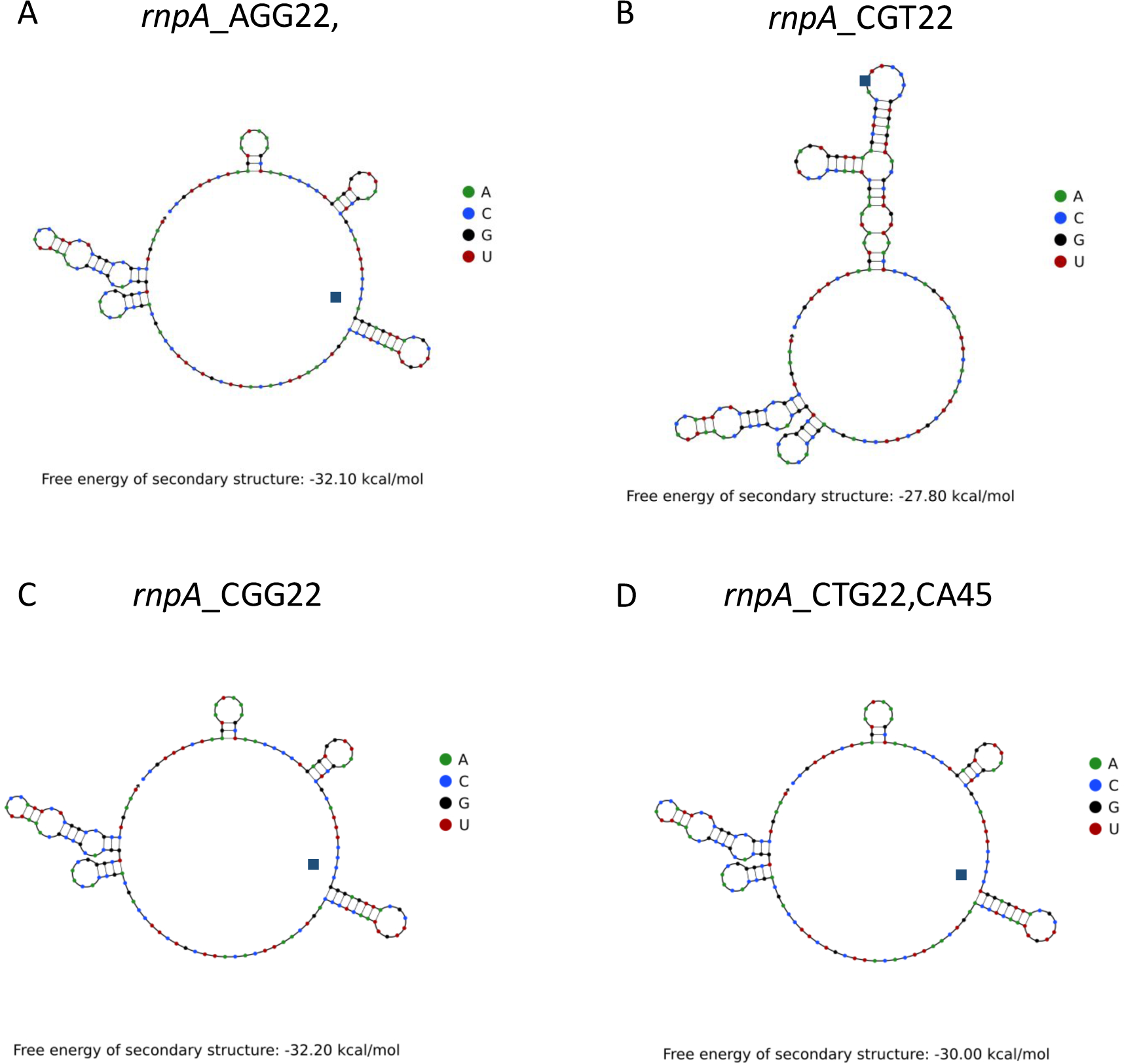
mRNA folding predictions for the gene *rnpA*. For folding predictions, we used 30 nucleotides upstream and100 nucleotides downstream of the rnpA start site using UNAfold (40). a) The wild-type *rnpA* sequence, with AGG (in blue box); b) The wild-type *rnpA* sequence with AGG→CGU in blue box (not observed); c) The wild-type *rnpA* sequence with AGG→CGG in blue box (observed with no growth rate defect); and d) The wild-type *rnpA* sequence with AGG→CTG in blue box and one complementary mutation CCC→CCA to maintain the mRNA loop (in blue box) (observed, also with no growth rate defect).

**Figure S6.**
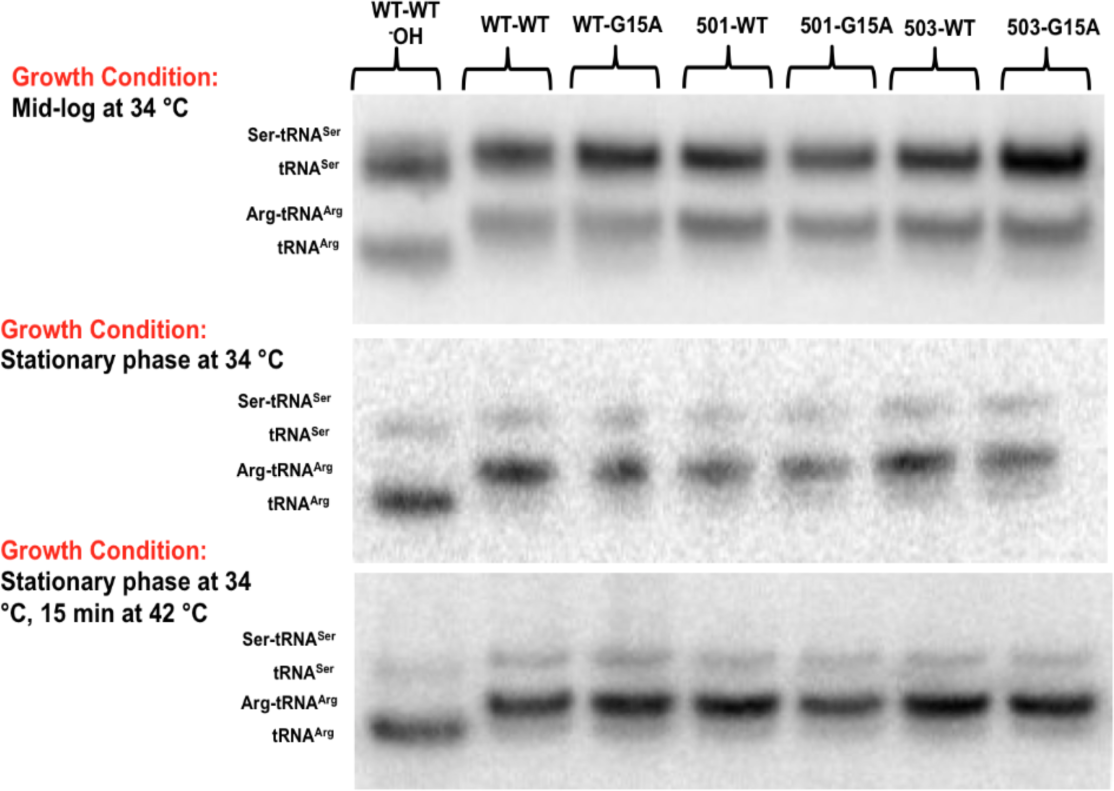
G15A ArgU does not affect expression and aminoacylation levels in WT and recoded *E. coli* strains. Northern blot Acid-Urea PAGE was performed on WT and G15A argU tRNA in wild-type *E. coli* (WT-WT and WT-G15A), and in the final strains C123a and b (501 and 503) at several growth conditions. Aminoacylation levels are comparable to wild-type for all conditions and combinations, suggesting no effect on charging levels despite the mutation sweeping into the population.

**Figure S7.**
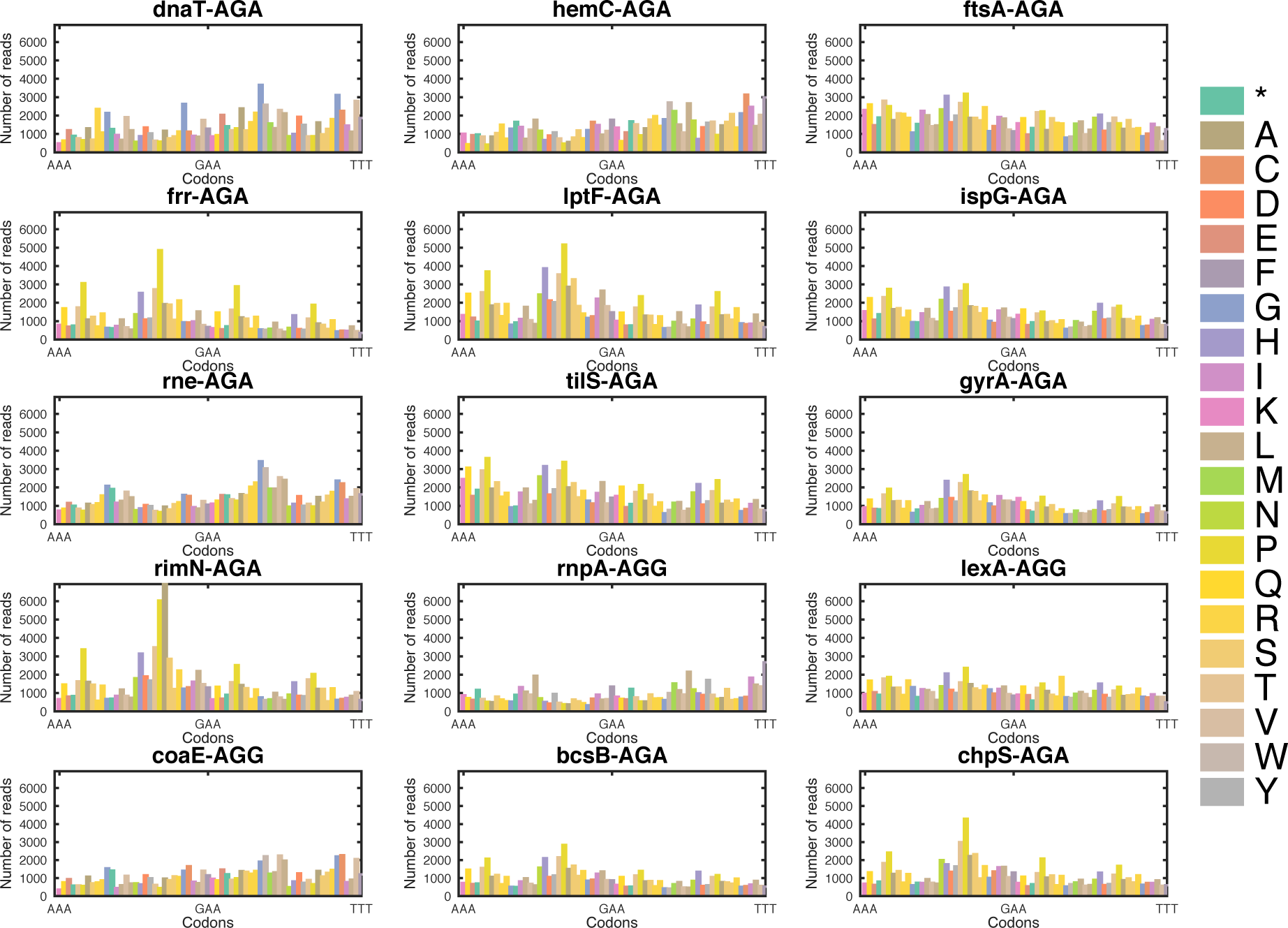
Number of reads for each codon and for each gene in the CRAM experiment at time point 24hrs. CRAM (Crispr-Assisted MAGE) was used to explore codon preference for several N-terminal AGR codons. The left y-axis (Number of reads) indicates abundance of a particular codon. The x-axis indicates the 64 possible codons ranked from AAA to TTT in alphabetical order. Experimental time point 24hrs is presented. Diversity was assayed by Illumina sequencing. Genes *bcsB* and *chpS* are non-essential and thus serve as controls for AGR codons that are not under essential gene pressure.

**Figure S8.**
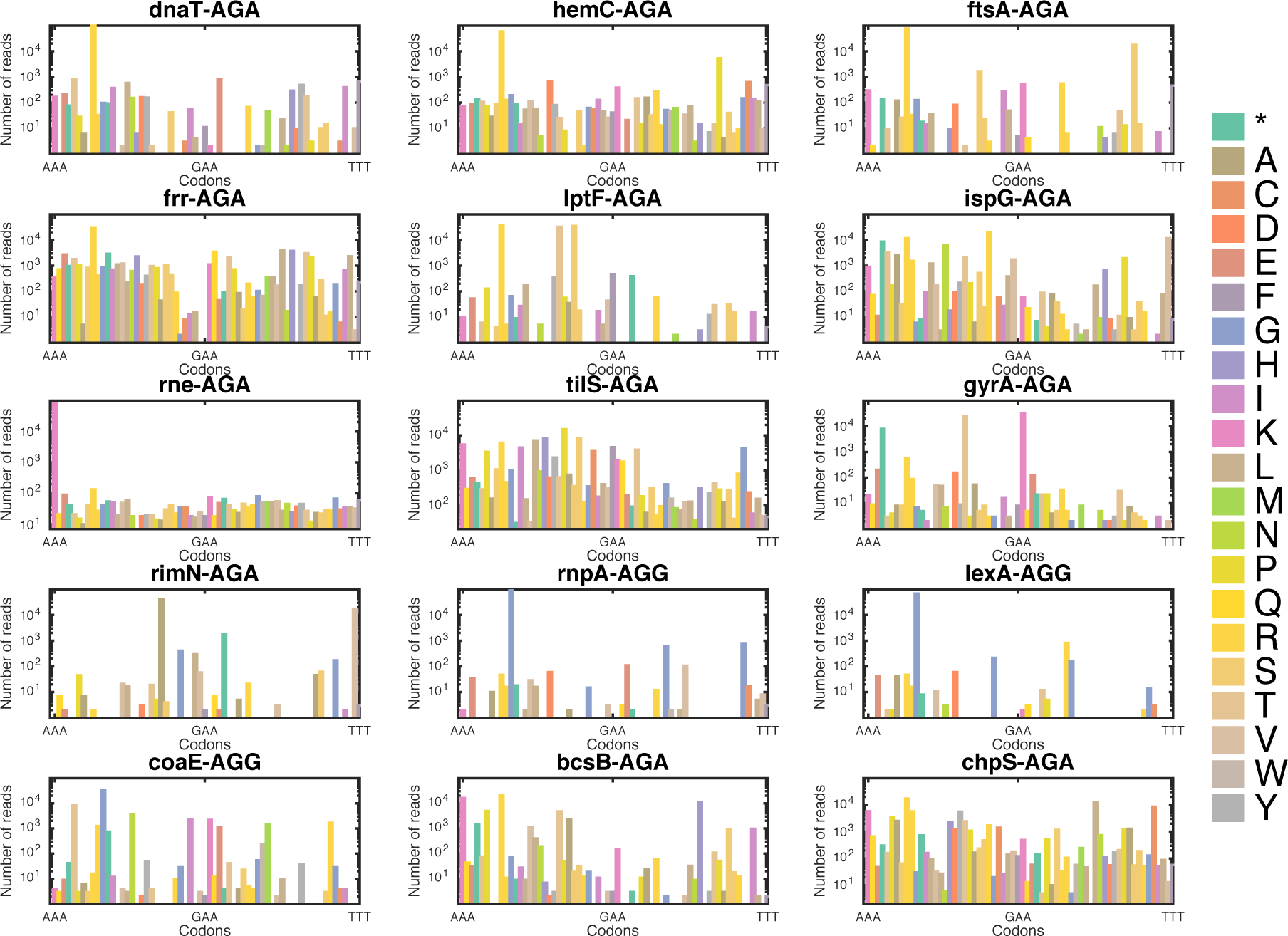
Number of reads for each codon and for each gene in the CRAM experiment at time point 144hrs. CRAM (Crispr-Assisted MAGE) was used to explore codon preference for several N-terminal AGR codons. The left y-axis (Number of reads) indicates abundance of a particular codon. The x-axis indicates the 64 possible codons ranked from AAA to TTT in alphabetical order. Experimental time point 144hrs is presented. Diversity was assayed by Illumina sequencing. Genes *bcsB* and *chpS* are non-essential and thus serve as controls for AGR codons that are not under essential gene pressure.

**Figure S9.**
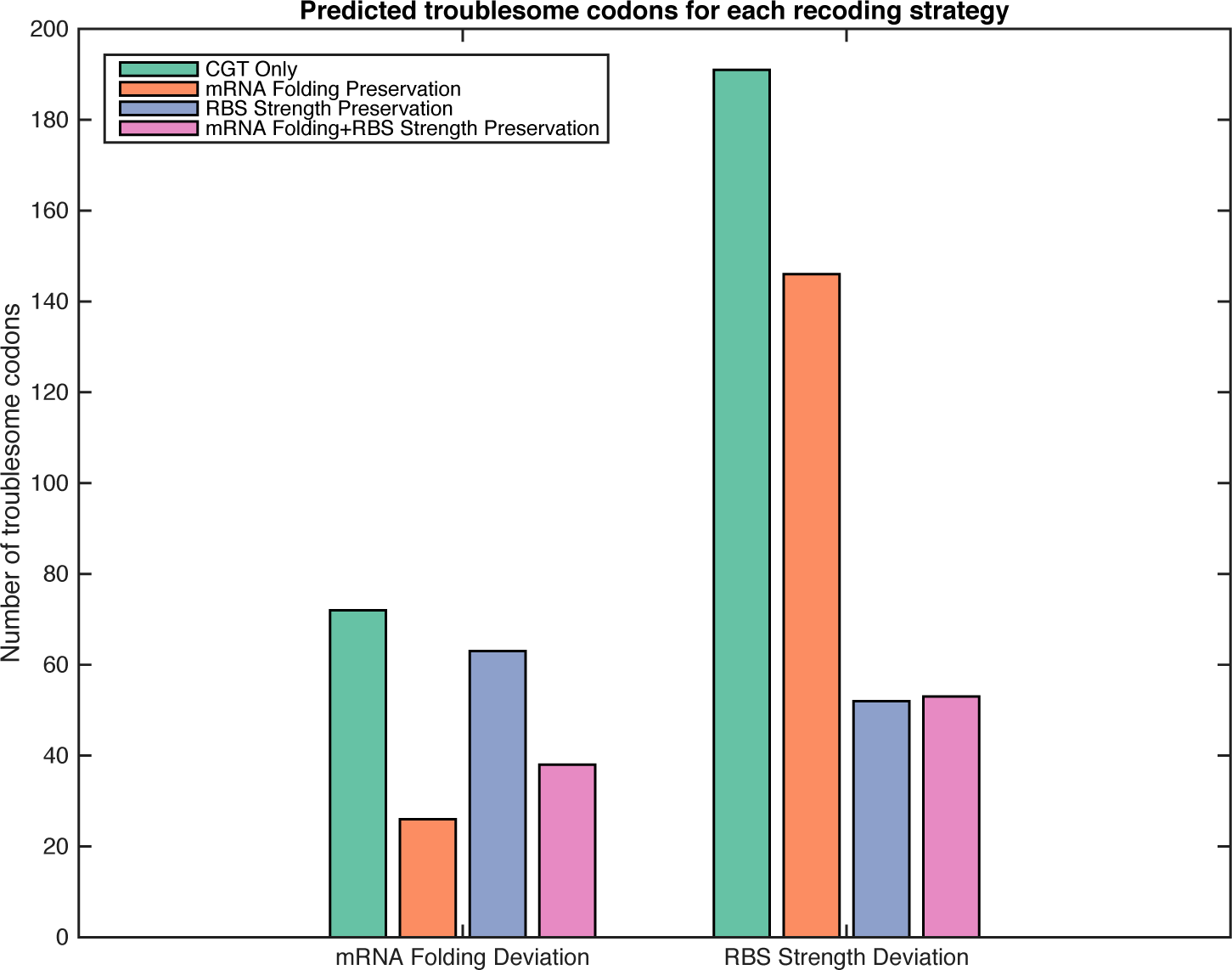
Number of predicted recalcitrant AGR codons for each AGR replacement strategy. 4 possible genomes replacing all 3222 AGRs have been designed using 4 replacement strategies. First AGRs were changed to CGU genome-wide (green bars). Second, AGR synonyms were chosen to minimize local mRNA folding deviation near the start of genes (orange bars). Third, AGR synonyms were chosen to reduce RBS strength deviation (blue bars). Finally, AGR synonyms were chosen to minimize both (purple bars). These genomes were then scored using a custom software available on Github (https://github.com/churchlab/agr_recoding), and compared. Every deviation outside of the Safe Replacement Zone is predicted to be a recalcitrant codon.

**Figure S10.**
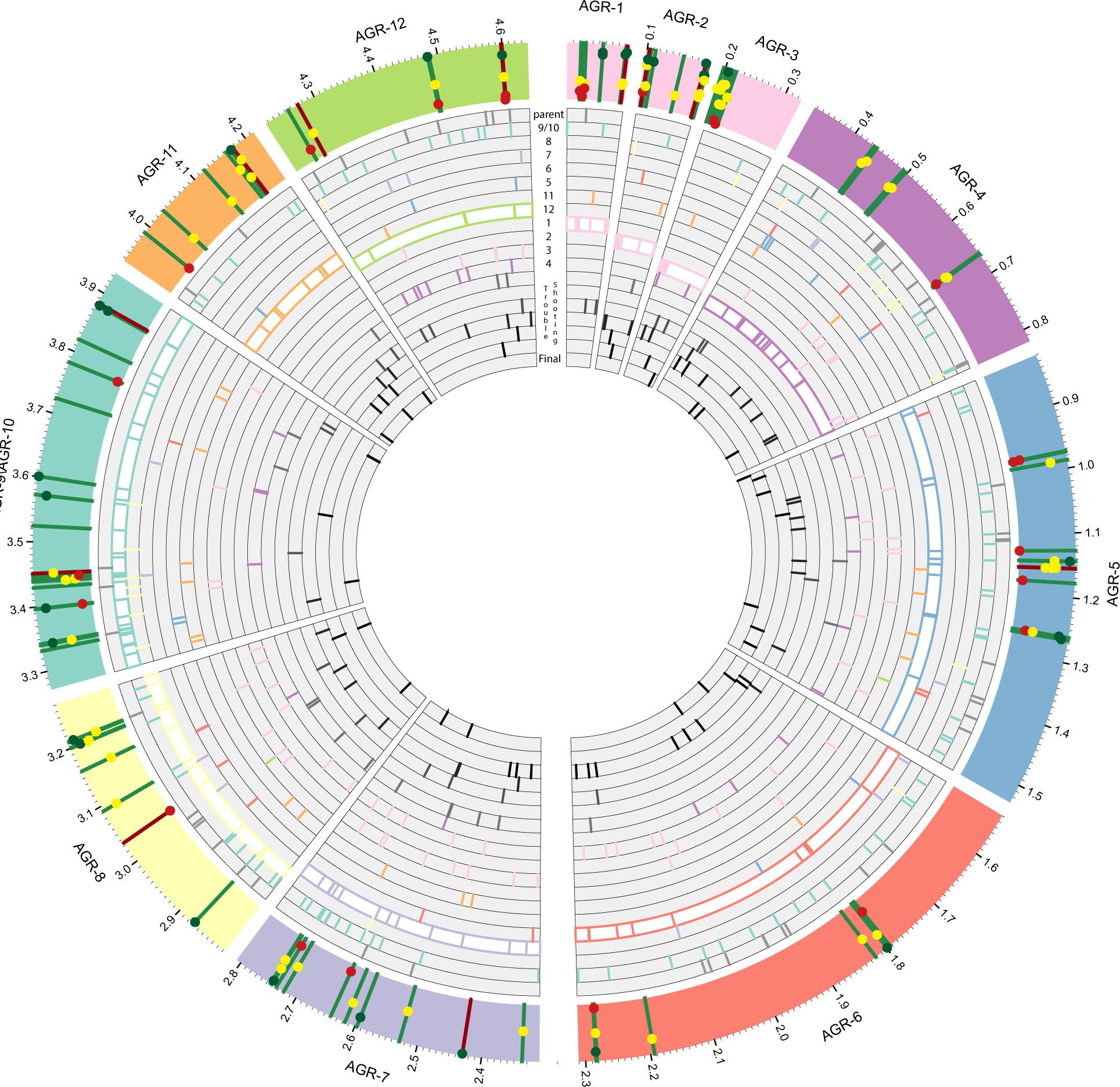
Representational graph of the fully recoded genome relative to MG1655. The outer ring contains the set grouping that each AGR codon (vertical line) is in. Each line contains information on troubleshooting (red if it required troubleshooting, green if not), and relative recombination frequency is represented by the position of the dot. Each internal ring represents the mutations that accumulated during strain construction. The target set of AGR codons for each ring is highlighted. The internal rings with black radial lines represent the mutations accumulated while the 13 recalcitrant codons were mutated to their optimized codon replacements.

